# The human placenta shapes the phenotype of decidual macrophages

**DOI:** 10.1101/2022.03.29.486171

**Authors:** Sigrid Vondra, Anna-Lena Höbler, Andreas Ian Lackner, Johanna Raffetseder, Zala Nikita Mihalic, Andrea Vogel, Leila Saleh, Victoria Kunihs, Peter Haslinger, Markus Wahrmann, Heinrich Husslein, Raimund Oberle, Julia Kargl, Sandra Haider, Paulina Latos, Gernot Schabbauer, Martin Knöfler, Jan Ernerudh, Jürgen Pollheimer

**Affiliations:** Department of Obstetrics and Gynecology, Reproductive Biology Unit, Maternal-fetal Immunology Group, Medical University of Vienna, Austria; Division of Inflammation and Infection (II), Department of Biomedical and Clinical Sciences (BKV), Linköping University, Linköping, Sweden; Otto Loewi Research Center, Division of Pharmacology, Medical University of Graz, Austria; Institute for Vascular Biology, Centre for Physiology and Pharmacology, Medical University Vienna, Austria; Christian Doppler Laboratory for Arginine Metabolism in Rheumatoid Arthritis and Multiple Sclerosis, Vienna, Austria; Department of Obstetrics and Gynecology, Reproductive Biology Unit, Placental Development Group, Medical University of Vienna, Austria; Division of Nephrology and Dialysis, Department of Medicine III, Medical University of Vienna, Vienna, Austria; Department of Obstetrics and Gynecology, Medical University of Vienna, Vienna, Austria; Center for Pathobiochemistry and Genetics, Institute of Medical Chemistry, Medical University of Vienna, Austria; Center for Anatomy and Cell Biology, Medical University of Vienna, Austria; Department of Clinical Immunology and Transfusion Medicine, Department of Biomedical and Clinical Sciences, Linköping University, Linköping, Sweden

**Author notes:** Correspondence to Jürgen Pollheimer. These authors contributed equally to this work.

## Abstract

During human pregnancy, placenta-derived extravillous trophoblasts (EVT) invade the decidua and communicate with maternal immune cells. The decidua can be distinguished into basalis (decB) and parietalis (decP), the latter being unaffected by placentation. By defining a novel gating strategy, we report accumulation of myeloid cells in decB. We identified a decidua basalis-associated macrophage (decBAM) population with a differential transcriptome and secretome when compared to decidua parietalis-associated macrophages (decPAMs). decBAMs are CD11c^hi^ and efficient inducers of Tregs, proliferate *in situ* and secrete high levels of CXCL1, CXCL5, M-CSF, and IL-10. In contrast, decPAMs exert a dendritic cell-like, motile phenotype characterized by induced expression of HLA class II molecules, enhanced phagocytosis, and the ability to activate T cells. Strikingly, EVT-conditioned media are able to convert decPAMs into a decBAM phenotype. Cumulatively, these findings assign distinct macrophage phenotypes to decidual areas depending on placentation and further highlight a critical role for EVTs in the induction of pregnancy-tolerant macrophage polarization.

**Graphical Abstract:** 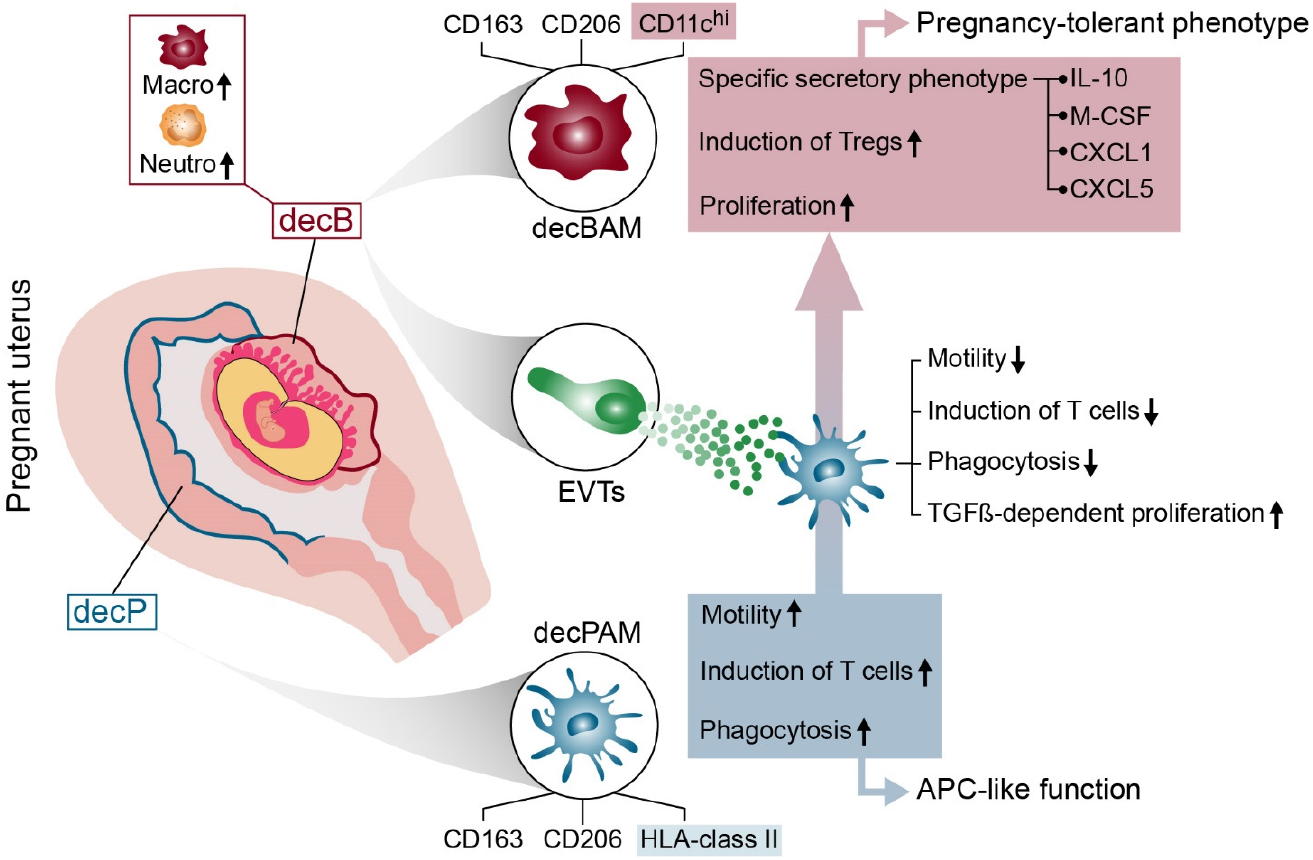

**Highlights:** In this study, we identified so far unrecognized, placenta-induced immune responses at the maternal-fetal interface. Altogether, we imply that placenta-derived trophoblasts induce a pregnancy-tolerant phenotype by suppressing antigen-presenting cell-like functions in maternal tissue macrophages.

## Introduction

During human pregnancy, fetal-derived placental extravillous trophoblasts (EVTs) invade the decidualized mucosal uterine surface, referred to as decidua (Knofler et al., 2019). This unique process, in which semi-allogeneic EVTs intermingle and communicate with maternal decidual immune cells, is fundamental for the maintenance and success of pregnancy. An imbalance in these interactions can lead to the development of obstetrical syndromes such as preeclampsia or uterine growth restriction (Brosens et al., 2011). The human decidua has been the subject of various studies directed towards understanding the function of uterine immune cell types. Under physiological conditions, the maternal-fetal interface is dominated by a non-cytotoxic immune cell response, which enables fetal tolerance and supports placental function (Erlebacher, 2013; Moffett et al., 2017). A central role in the maintenance of this tissue-specific, pregnancy-sustaining environment has been assigned to an interactive network between EVTs and maternal immune cells. Remarkably, decidual natural killer (dNK) cells and macrophages cooperate to transform the decidual vasculature into dilated conduits able to deliver high amounts of low-pressure blood to the growing fetus (Pollheimer et al., 2018). Genetic data suggest that the specific combination of killer-cell immunoglobulin-like receptors expressed on dNK cells and their MHC ligands on EVTs are crucial in determining reproductive success (Hiby et al., 2010). Moreover, there are also indications for an active suppression of immune responses at the maternal-fetal interface. In humans, EVTs have been demonstrated to induce a non-cytotoxic phenotype in dNK cells by initiating intercellular trafficking of the non-classical MHC-I class molecule HLA-G (Tilburgs et al., 2015). In addition, studies in mice suggest an active blockage of T cell infiltration at the maternal-fetal interface even after forced reactivation of maternal memory T cells by placenta-specific overexpression of ovalbumin (Nancy et al., 2012). However, little is known about the specific function of decidual macrophages. A previous paper reported two distinct macrophage populations expressing either high or low levels of cell surface CD11c. The authors further suggested that the CD11c^lo^ population comprise an antigen-presenting cell (APC) population (Houser et al., 2011). Recently, a single-cell transcriptomic approach identified various maternal macrophage subpopulations in the human decidua, two of which constitute tissue resident cells, referred to as dM1 and dM2, and one is found in the intervillous space adhering to placental villi, designated as dM3 (Vento-Tormo et al., 2018). The dM3 type was recently characterized and identified as placenta-associated maternal macrophages (PAMMs) (Thomas et al., 2021). In this paper, the authors also unraveled distinctive phenotypes of fetal-derived placental macrophages, referred to as Hofbauer cells (HB). Different to HBs and PAMMs in the intervillous space, decidual tissue-resident macrophages are not uniformly confronted with placental tissues. After the blastocyst becomes embedded in the endometrium and the placenta develops, the formation of the maternal-fetal interface occurs only at a specific uterine site, the decidua basalis (decB). In contrast, large areas of the decidua, the decidua parietalis (decP), are unaffected by placentation. While various morphological differences such as degree of vascularization (Plaisier et al., 2007; Windsperger et al., 2020) or tissue stiffness (Abbas et al., 2019) have been noticed, little is known about the difference between decB and decP tissues in terms of immune cell content and activation. In a recent publication, we were able to show low abundance of T cells in decB tissues when compared to decP samples. The drop in both CD4^+^ helper and CD8^+^ cytotoxic T cells was further associated with low abundance of T cell-recruiting high endothelial venules (Windsperger *et al*., 2020). These findings imply fundamental differences in immune cell distribution and extravasation into decB and decP. However, to date, there are no comparative data on immune cell distribution or function in decB and decP tissues that allow conclusions to be drawn about placentation-dependent effects on decidual immune cells.

In this study, we set out to select donor-matched first-trimester decB and decP tissue samples in order to identify immunological adaptation that specifically occur in response to placentation. By defining a novel flow cytometry gating strategy, we demonstrate that tissue-resident macrophages and neutrophils accumulate in decB tissues. We further identified a specific phenotype and functionality of decB-associated macrophages (decBAMs) in comparison to decP-associated macrophages (decPAMs). While decBAMs show a higher capacity to proliferate and induce Treg formation, decPAMs are more motile and phagocytic, and efficiently induce T cell proliferation.

## Results

### Myeloid cells specifically accumulate at the maternal-fetal interface

In this study, we aimed at identifying placentation-specific immune modulation in the human decidua during early human pregnancy. To this end, we established a cohort of first-trimester decidual samples including areas affected by placentation (decidua basalis, decB) and donor-matched areas that do not interact with villous placental tissues (decidua parietalis, decP) (Fig. 1A). The absence or presence of HLA-G, a marker for EVTs, served as an additional indicator to distinguish decP from decB (Fig. 1, B and C), respectively. An overview of our tissue sampling strategy is outlined in Fig. S1, A and B. First, decB and decP samples were morphologically identified based on well-described characteristics (Loke, 1996). Decidua parietalis tissue has a visible smooth mucosal covering (Fig. S1 B), shows the presence of glands and luminal epithelium (Fig. S1 C) as well as a significantly higher cellular density when compared to matched decB samples (Fig. S1 D). In contrast, we identified decB by its hemorrhagic appearance and the presence of numerous EVTs invading the stroma and vasculature (Fig. S1 B and Fig. S1 E). Overall, we found that cell isolates from first-trimester decB tissues contain on average 11 % (± 4.34 % S.D.) of HLA-G^+^ EVTs compared with < 0.05 % of EVTs in decP tissues (Fig. 1 D) and that the total cellular density is significantly lower in decB when compared to matched decP samples (Fig. S1 C). Next, we defined the distribution of the major immune cell subtypes in decB and decP tissues, revealing an accumulation of myeloid CD45^+^CD14^+^ monocytes/macrophages and CD45^+^CD66b^+^ neutrophils in decB samples (Fig. 1 E). A representative flow cytometry gating strategy is shown Fig. S2 A and B. We also considered contaminating blood a possible reason for the decB-specific neutrophil accumulation. To this end, we examined decB tissues by H&E and IF stainings against CD66b and confirmed the presence of interstitial neutrophils (Fig. S1 E). Ficoll^®^ gradient centrifugation revealed a significantly higher CD66b^+^ low-density neutrophil population in decB (mean = 41.48 %, ± 14.57 % S.D., n = 4) when compared to decP (mean = 3.37 %, ± 1.08 % S.D., n = 4) or blood samples from pregnant (mean = 1.29 %, ± 0. 19 % S.D., n = 3) and non-pregnant women (mean = 0.18 %, ± 0.13 % S.D., n = 3) (Fig 1 F). Moreover, while decB-derived CD66b^+^ neutrophils show a typical CD14^-^, HLA-DR^-^, CD15^dim^, CD11c^dim^, CD33^dim^, CD11b^+^ and CD16^+^ expression pattern, they lack the blood neutrophil marker CD62L (L-selectin) (Fig. S1 F). As our focus in this study was on decidual macrophages, we re-calculated our established immune cell distribution by excluding CD66b+ neutrophils, resulting in a significantly higher proportion of macrophages in decB tissue isolates (Fig. S1 G). Well in line, comparative analyses of secreted factors from decB and decP tissue explants revealed an overrepresented myeloid-associated cytokine signature in decB tissues, including IL-10, CXCL1, CCL3, CCL4, CXCL5, and suppressors of T cell responses (TRAIL and PD-L1). In contrast, decP samples showed elevated levels of factors involved in T cell, DC, and NK cell immunity such as IL-7, CXCL1, sCD5, sCD6, TRANCE or Flt3 (Fig. 1 G).

**Figure 1.**
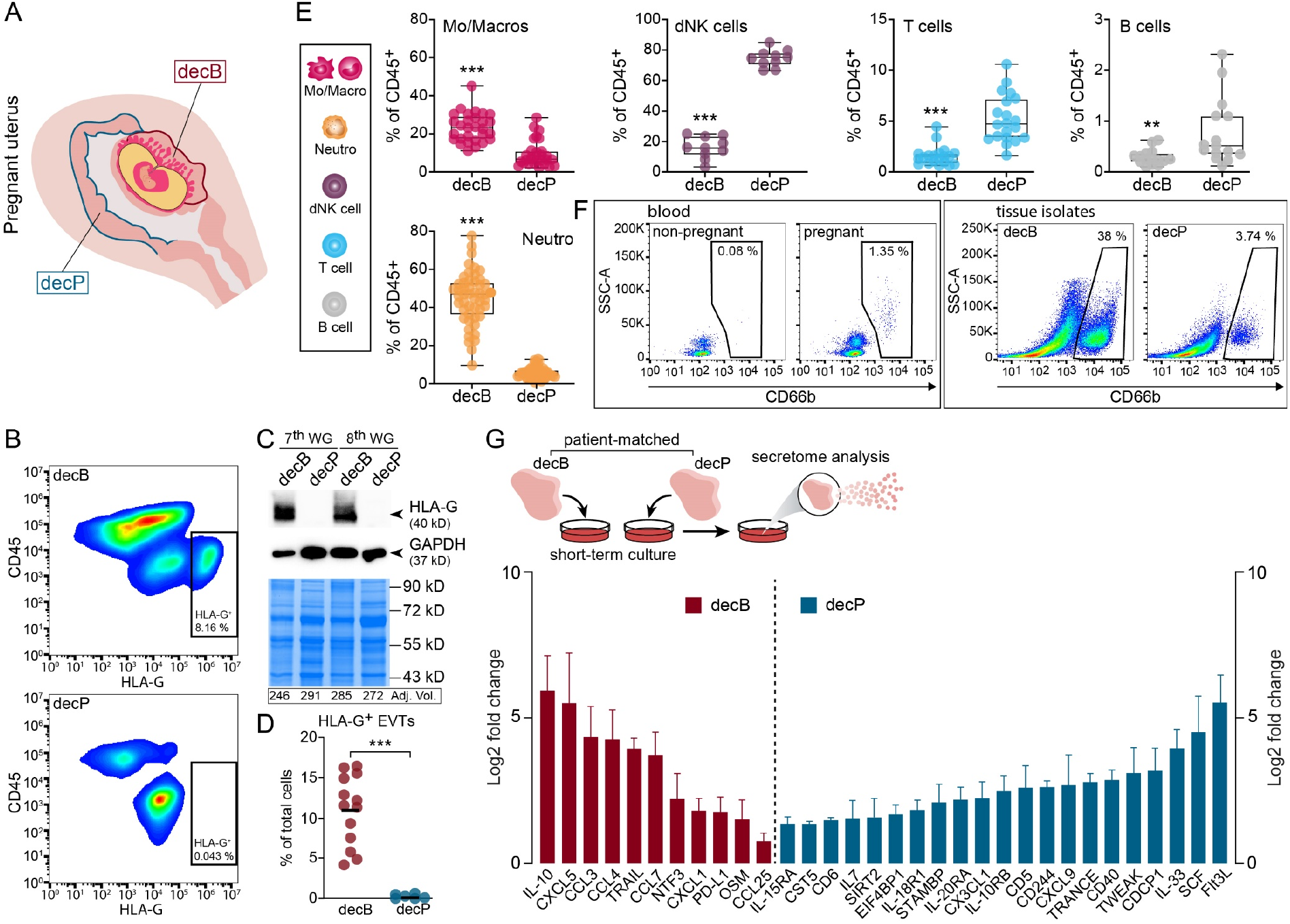
Investigation of immune cell distributions in first-trimester decidual samples identifies myeloid cell accumulation in first-trimester decB tissues. **(A)** Schematic drawing of a human pregnant uterus during the first trimester of pregnancy indicating decidua basalis (encircled by a red line, decB) and decidua parietalis (encircled by a blue line, decP). **(B)** Representative flow cytometry plots of decB and decP tissue cell isolates demonstrating distribution of CD45^+^ immune cells and HLA-G^+^ EVTs. The percentages of HLA-G^+^ EVTs are indicated. **(C)** A representative western blot of two sets of donor-matched decB and decP protein lysates probed with antibodies against HLA-G is shown. The expression of GAPDH was determined to control for successful protein transfer and immunodetection. A stain-free system guaranteed equal total protein loading prior to blotting. Adjusted volumes (Adj. Vol.) of total band intensities are indicated underneath the stain-free gel images. **(D)** Quantification of HLA-G^+^ EVTs in first-trimester decB (n = 14) and decP (n = 7) cell isolates, analyzed by Student’s t-test. The center line represents the mean. **(E)** Bar graphs show the percentages of Mo/Macros (n = 31 pairs), dNK cells (n = 13 pairs), T cells (n = 21 pairs), B cells (n = 16 pairs) and Neutros (n = 53 pairs) of the total CD45^+^ leukocyte population in donor-matched decB and decP tissue isolates. **(F)** Representative flow cytometry plots of isolated PBMCs from the blood of healthy non-pregnant and pregnant female donors (left panel) and donor-matched decB and decP tissue isolates (right panel). The gates represent the frequency of low-density CD66b^+^ neutrophils of the CD45^+^ PBMC fraction obtained by LymphoPrep®. **(G)** Summary data (log2 fold change, n = 4) for the significant differences in secreted factors from patient-matched decB and decP tissue explants after 20 h in culture measured by the proximity extension assay (www.Olink.com). The schematic drawing shown on the top illustrates the experimental set-up. Data are represented as mean + SD (standard deviation). Significances were calculated using multiple t-tests. **, P ≤ 0.01. decGE, decidual glandular epithelium; EVT, extravillous trophoblast; Mo/Macro, monocytes, macrophages; Neutro, neutrophils; dNK cell, decidual natural killer cell; WG, week of gestation.

### Formation of decB is associated with the accumulation of tissue-resident macrophages

To further increase the resolution of our data we applied specific markers to dissect macrophages from infiltrating blood monocytes. *In situ* double and triple immunofluorescence (IF) staining confirmed that all CD14^+^ cells residing in the decB and decP are positive for the pan macrophage marker CD68 and double-positive for CD163 and CD206 (Fig. S3 A - C). Subsequent flow cytometry analysis of decidual cell isolates revealed two CD14^+^ populations, one lacking CD163 and CD206 expression and the other highly positive for both tissue macrophage markers (Fig. 2 A). While the latter CD163^+^CD206^+^ population showed a typical enlarged cell size and morphology with abundant vacuoles as well as high autofluorescence, as is commonly observed in macrophages (McGovern et al., 2014), CD14^+^CD163^-^CD206^-^ cells displayed a morphology typical for monocytes such as smaller cell size or little granularity (Fig. 2, B and C). Bulk RNA-seq confirmed this notion by demonstrating that CD14^+^CD163^+^CD206^+^ cells express typical hallmark genes of tissue-resident macrophages such as *MERTK, CD68, MAFB* or *FCGR2A* (Xue et al., 2014) and express comparatively low levels of monocyte marker genes including L-selectin (*SELL*), *S100A12, S100A8*, and *S100A9* (Fig. 2 D and Fig. S3 D). Indeed, IF-analysis shows that CD14^+^S100A12^+^ monocytes are found in the lumen of decidual vessels and not in the tissue stroma (Fig. S3 E).

**Figure 2.**
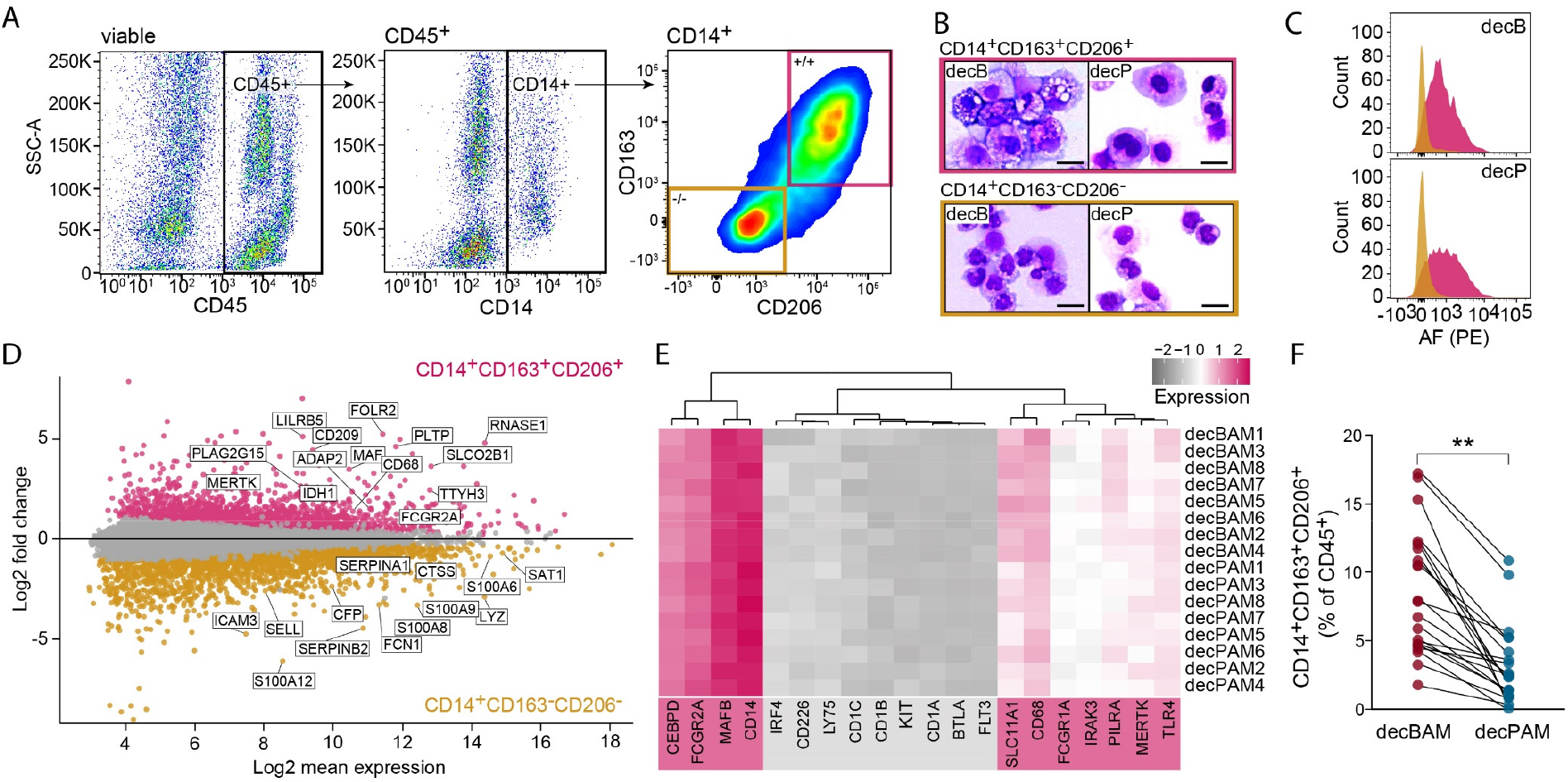
Isolation and characterization of tissue-resident decidual macrophages. **(A)** Flow cytometry gating strategy used to identify tissue-resident decidual macrophages by a CD45^+^CD14^+^CD163^+^CD206^+^ expression profile. Pink gate, CD14^+^CD163^+^CD206^+^ macrophages. Orange gate, CD14^+^CD163^-^CD206^-^ monocytes. **(B)** Macrophage (pink rectangle) and monocytic (orange rectangle) populations from decB and decP were sorted and stained by Giemsa staining. Representative images of *n* = 4 experiments. Scale bar, 20 µm. **(C)** Representative flow cytometry plots identifying autofluorescence (AF) in isolated CD14^+^CD163^+^CD206^+^ (pink histogram) and CD14^+^CD163^-^CD206^-^ (orange histogram) cell isolates from decB and decP. **(D)** MA plot of differentially expressed genes (DEGs) from RNA-seq. Pink and orange dots represent significantly up- or down-regulated DEGs in isolated CD14^+^CD163^+^CD206^+^ (n = 8 pairs) and CD14^+^CD163^-^CD206^-^ (n = 4 pairs) cells from decB and decP tissues. **(E)** Heatmap showing expression of macrophage (underlaid with pink)- and DC (underlaid with grey)-associated hallmark genes in isolated decBAM and decPAM. **(F)** Dotplot demonstrating distribution of CD14^+^CD163^+^CD206^+^ decBAMs and decPAMs in patient-matched first-trimester decB and decP samples (n = 21 pairs). Patient-matched samples are indicated via connected dot plots. P-value was generated using paired t-test. **, P ≤ 0.01. decB, decidua basalis; decP, decidua parietalis; decBAM, decB-associated macrophage; decPAM, decP-associated macrophage.

Next, we analyzed whether our isolated decidual macrophage populations may express DC core markers retrieved from various hallmark papers deciphering macrophage and DC signature gene expression in human and mice (Gautier et al., 2012; McGovern *et al*., 2014; Tamoutounour et al., 2013). As expected, decidual macrophages express conserved lineage markers such as *MERTK* and *MAFB* as well as other typical markers for human and mouse macrophages, including *CD14, CD68, CEBPD, FCGR1A* (CD64) or *FCGR2A* (CD32). In contrast, DC-associated lineage markers such as *CD1A*/*B*/*C, CD226, FLT3* or *KIT* were absent from both populations (Fig. 2 E). By considering our established markers for tissue-residency we were able to show a decB-associated accumulation of tissue macrophages (Fig. 2 F). Finally, we termed macrophages obtained from different decidual compartments decBAMs and decPAMs, respectively. These data show that placentation is associated with increased numbers of true tissue-resident macrophages.

### Accumulation of decidual tissue macrophages is associated with *in situ* proliferation

We next sought to determine the potential of decB to recruit neutrophils and macrophages. Therefore, we generated conditioned media (CM) from cultivated patient-matched decB and decP tissue samples and performed chemoattractant assays using isolated PBMCs or neutrophils from peripheral blood samples. These assays revealed no differential effect between decB- and decP-CM in the recruitment of CD14^+^ blood-derived monocytes (Fig. 3 A) or other PBMCs, including T cell subtypes, B cells or NK cells (Fig. 3 A, Fig. S4 A). In contrast, separate assays showed that decB-CM exert a significantly higher potential to attract blood neutrophils (Fig. 3 B). Well in line, culture media of decB tissues contained high levels of well-described neutrophil chemoattractants including CXCL1, CXCL8, S100A8, and S100A9 (Kolaczkowska and Kubes, 2013; Zenz et al., 2005) (Fig. 3 C). Strikingly, *in situ* IF staining showed significantly more CD14^+^Ki67^+^ macrophages in decB tissues when compared with decP sections (Fig. 3 D). Decidual tissue macrophages also expressed a wide array of cell cycle markers including CCNA1, pRB, and CDK1 (Fig. 3 E). In addition, we observed the presence of mitotic figures in tissue-resident CD14^+^ decidual macrophages (Fig. 3 E). Flow cytometry-sorted decB macrophages Finally, an *ex vivo* proliferation assay determining EdU incorporation into cultivated decidual explants revealed that decBAMs are capable of *de novo* DNA synthesis, again to a greater extent than decPAMs (Fig. 3 F). Representative low magnification pictures showing CD14^+^ ki67^+^, CD14^+^CCNA1^+^, CD14^+^ CDK1^+^, CD14^+^pRB^+^, and CD14^+^ EdU^+^ decBAMs are represented in Fig. S4 B - D.

**Figure 3.**
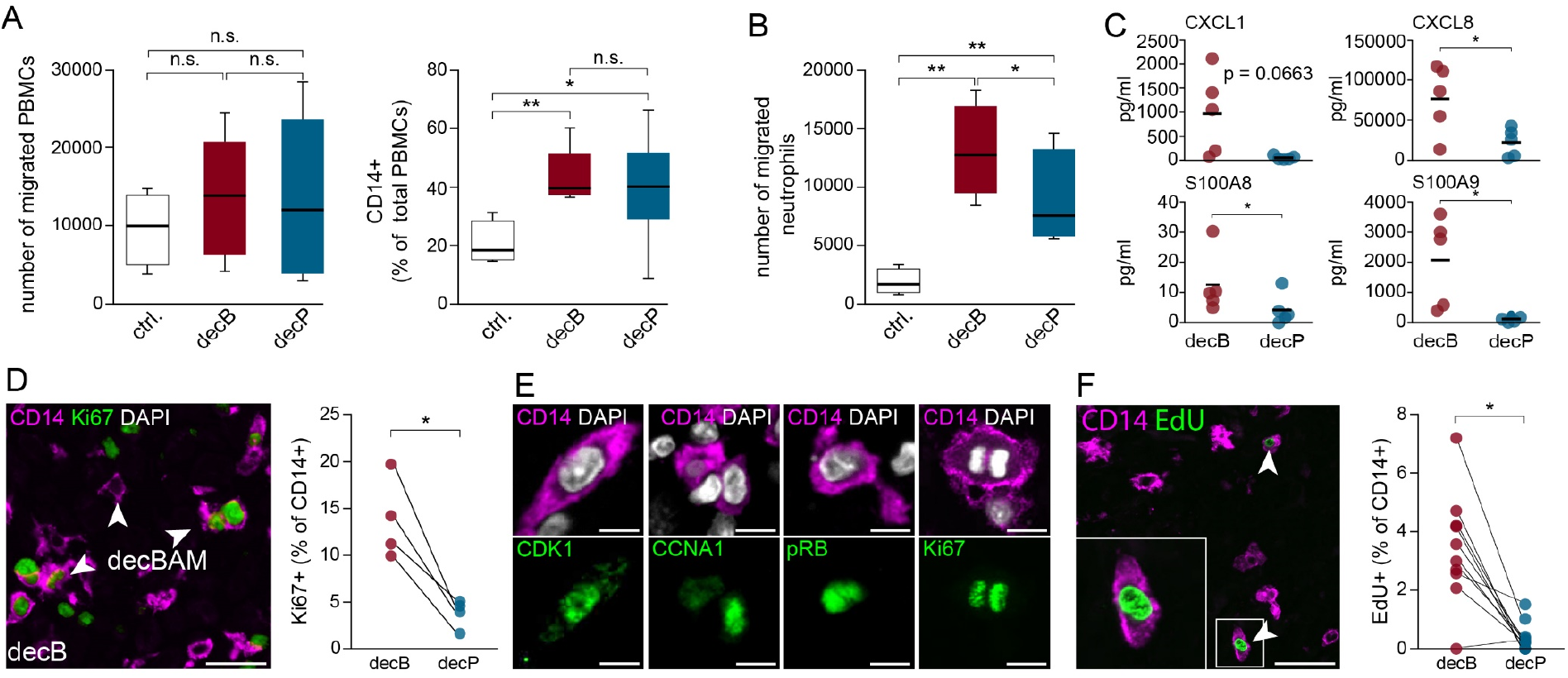
decB-associated accumulation of myeloid cells is accomplished by chemoattractant factors and local proliferation of decBAMs. **(A)** Migratory response of PBMCs (left diagram) and CD14^+^ blood cells (right diagram) to culture medium containing decB- or decP-CM (1:4, n = 8). Decidua culture medium was used as a control. P-values were generated using Repeated-measures ANOVA with Tukey’s multiple comparison test. **(B)** Migratory response of blood neutrophils to culture medium without or with decB- and decP-CM (1:2, n = 4). Boxes in box plots represent upper and lower quartiles. The center line represents the median. Whiskers show the minimum and maximum of the data. P-values were generated using Repeated-measures ANOVA with Tukey’s multiple comparison test. **(C)** Secretory profile of cultivated (20 hours), patient-matched decB and decP explants (n = 5 pairs) measured by bead-based multiplex immunoassays. Dots show measured levels in pg/ml. The center line represents the mean. P-values were generated using paired t-tests. **(D)** Representative IF staining of a first-trimester decB tissue section showing CD14 (magenta) and Ki67 co-staining (green). Quantification of four patient-matched decB and decP tissue samples is shown to the right by a dot plot. Each dot represents the percentage of Ki67^+^CD14^+^ double-positive cells for each tissue sample (n = 4 pairs). Scale bar, 20 µm. Patient-matched samples are shown in connected dot plots. P-value was generated using a paired t-test. **(E)** IF co-staining of first-trimester decB tissue sections showing cell cycle markers including CDK1, CCNA1, pRB in green and CD14 in magenta. The two images to the right upper and lower panel demonstrate a Ki67 expressing mitotic figure in a CD14^+^ cell. DAPI (white) was used to visualize nuclei. Scale bars, 2 µm. **(F)** Representative IF staining of a cultivated decB explant tissue demonstrating EdU (green) incorporation into CD14^+^ (magenta) macrophages, indicated by arrowheads. Inset picture shows a EdU+ macrophage at higher magnification. Quantification of EdU incorporation into CD14^+^ macrophages of cultivated decB and decP first-trimester tissue explants. Each dot represents the percentage of CD14^+^ EdU^+^ cells for each tissue sample evaluated (n = 10 pairs). Patient-matched samples are indicated via connected dot plots. Scale bar, 20 µm. P-value was generated using a paired t-test. *, P ≤ 0.05; **, P ≤ 0.01, n.s.: not significant. decB, decidua basalis; decP, decidua parietalis.

These results suggest that decB-specific accumulation of tissue macrophages is driven by *in situ* proliferation and that, as expected, neutrophils are specifically recruited from blood.

### decBAMs show a distinctive transcriptional and secretory phenotype

In-depth analysis of our bulk RNA-seq-based transcriptome data revealed substantial differences between isolated patient-matched decBAMs and decPAMs, which are shown to form two distinct groups by principal component analysis (PCA) (Fig. S5 A). We identified 1127 significantly differentially expressed genes (DEGs, adjusted p value < 0.05) between decBAMs (512 up) and decPAMs (615 up) (Fig. 4 A). Immune-related DEGs included *ITGAX* (CD11c), *CD44, CXCL2*, and *ANPEP* in decBAMs, and *AXL, GAS6, SELENOP, C3*, and *IL2RA* in decPAMs (Fig. 4 A). IF- and flow cytometry-based analyses confirmed higher expression of HMOX1, CD44, and CD11c protein in decBAMs (Fig. 4 B). Single channels and lower magnification pictures of CD44 and HMOX1 stainings are shown in Fig. S5 B and C. Moreover, functional gene enrichment analysis revealed that gene sets involved in granulocyte/neutrophil activation and myeloid immune responses were overrepresented in decBAM-associated DEGs (Fig. S5 D). CD11c has already been suggested as a marker to distinguish between different decidual macrophage populations, however not in the context of regionally distinct decidual tissues (Houser *et al*., 2011). DEGs highly expressed in CD11c^hi^ macrophages identified by Houser et al. mainly coincide with decBAM upregulated genes, and conversely, CD11c^lo^ associated transcripts show a clear overlap with decPAM-associated gene signatures (Fig. S5 E). Prior to our study, three tissue-resident myeloid cell types have been identified by single-cell (sc) RNA-seq, designated as decidual macrophage 1 (dM1), decidual macrophage 2 (dM2) and dendritic cell 1 (DC1) (Vento-Tormo *et al*., 2018). A third decidual macrophage subtype (dM3) identified in this study, later defined as a subtype of placenta-associated maternal macrophages (PAMMs), does not constitute a decidua-resident cell type but is found in the intervillous blood space, adhering to placental villi (Thomas *et al*., 2021). Our comparative computational approach showed that transcripts upregulated in decPAMs clearly associated with the dM2 phenotype whereas decBAMs showed a strong overlap with the previously identified dM1-associated genes both in global and single gene expression pattern (Fig. 4 D). Well in line, decBAM- or decPAM-associated DEGs identified by our bulk RNA-seq analysis were also differentially regulated in dM1 and dM2 populations (Fig. 4 E). In addition, decBAMs showed a closer similarity with dM3-specific signatures, compared to decPAM (Fig. S5 F). To this end, we isolated PAMMs and confirmed their transcriptional overlap with dM3 signatures but also with maternal blood-derived immune cells such as monocytes or dendritic cells (DC2) (Fig. S 5 F). Overall, these data confirm that PAMMs likely originate from blood monocytes (Thomas *et al*., 2021). Next, we digested decB and decP tissues in order to study the secretome of isolated and short-term cultivated EVTs, neutrophils, decBAMs, and decPAMs (Fig. 4 F). All cells were sorted by FACS except for EVTs, which were isolated via magnetic beads conjugated with anti-HLA-G. Due to their large size, EVTs are very sensitive to the high pressure and shear forces in a cell sorter and show improved viability when sorted over magnetic columns. Viability and purity of EVTs were confirmed by flow cytometry after isolation (Fig. S5 G).

**Figure 4.**
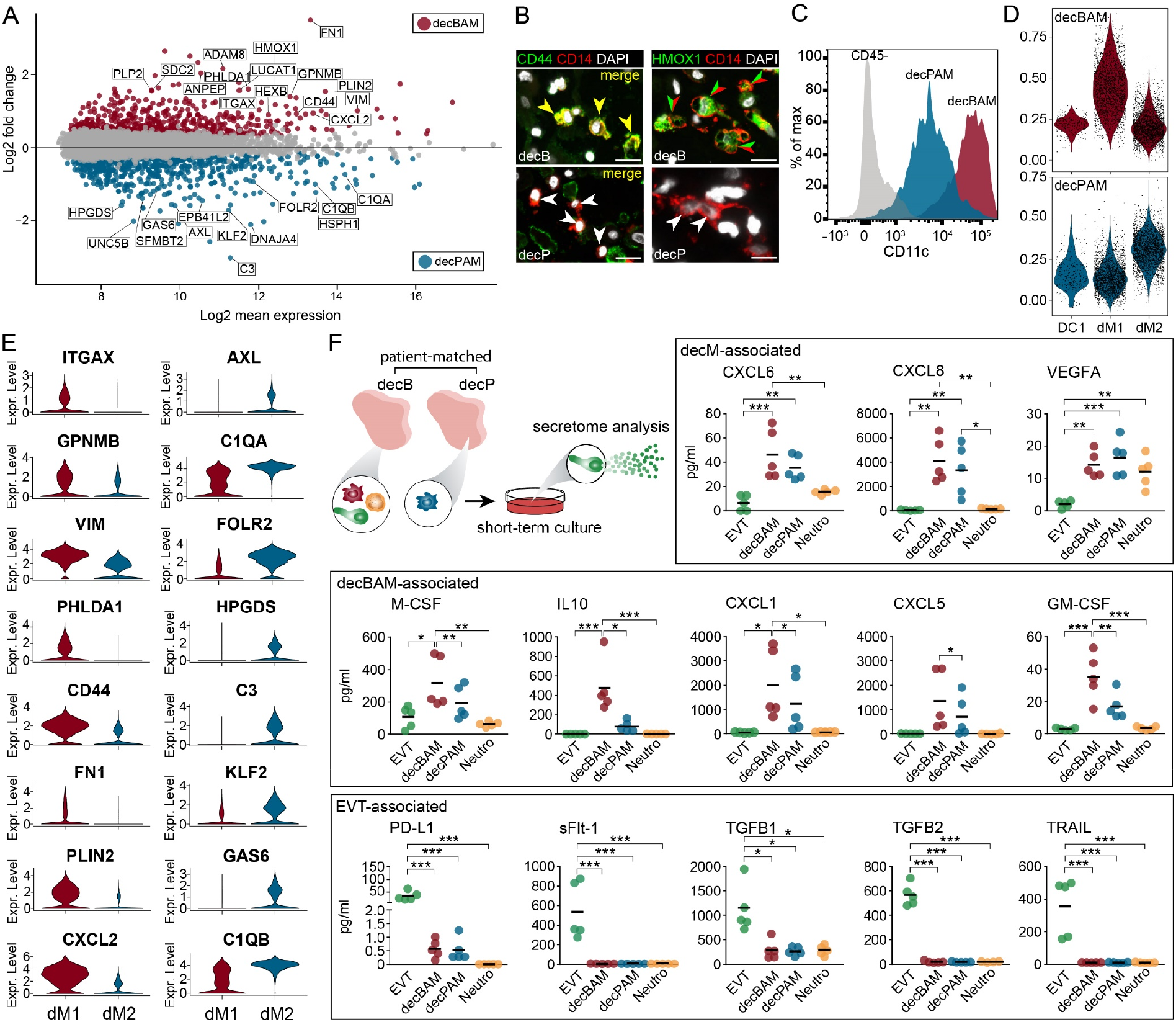
First-trimester decBAMs and decPAMs differ in their transcriptional and secretory profile. **(A)** MA plot of DEGs from RNA-seq. Red and blue dots represent significantly up- or down-regulated DEGs in patient-matched isolated CD14^+^CD163^+^CD206^+^ decBAMs and decPAMs (n = 8 pairs). **(B)** Representative IF co-staining of a first-trimester paraffin-embedded decB and decP tissue section showing HMOX1 and CD44 expression (green) in CD14^+^ tissue macrophages (red). Arrowheads in yellow (left) or in green and red (right) indicate double positive cells. Scale bars, 20 µm. **(C)** Representative flow cytometric analysis of isolated decBAMs and decPAMs showing CD11c expression. **(D)** Violin plots showing expression of decBAMs (red) and decPAMs (blue) signatures in selected scRNA-seq clusters (Vento-Tormo et al., 2018). **(E)** Violin plots presenting expression of selected decBAM- and decPAM-associated gene transcripts in dM1 and dM2 scRNA-seq clusters (Vento-Tormo et al., 2018). **(F)** Secretory profile of magnetically sorted and cultivated (20 h) EVTs, decBAMs, decPAMS and neutrophils, determined by bead-based multiplex immunoassays (n = 5 pairs). Measured levels (pg/ml) of growth factors, chemokines and cytokines are shown as dots and the group mean is shown as centered line. The schematic drawing shown on the top illustrates the experimental set-up. P-values were generated using One-way ANOVA with Tukey’s multiple comparison test. *, P ≤ 0.05; **, P ≤ 0.01; ***, P ≤ 0.001. decB, decidua basalis; decP, decidua parietalis; decM, decidual macrophage; decBAM, decB-associated macrophage; decPAM, decP-associated macrophage; EVT, extravillous trophoblast; Neutro, neutrophil.

We determined secretion of cytokines and growth factors, which were selected based on our previous analyses and their involvement in myeloid or neutrophil immunity and function. In parallel, we measured these factors in decB and decP tissue explant CMs to control for a possible loss of tissue specificity upon single cell culture (Fig. S6 A). While decBAMs culture media contained significantly higher levels of CXCL1, CXCL5, GM-CSF, M-CSF, and IL-10, decBAMs and decPAMs were equally efficient in releasing CXCL6, as well as high levels of CXCL8. Among the macrophage-associated factors, the measured GM-CSF levels were the lowest. All three immune cell populations secreted low but significantly induced levels of VEGFA, when compared to EVTs. Our analyses further revealed that EVTs are efficient producers of sPD-L1, TGFβ1, TGFβ2 and TRAIL. As a control for cell identity and differentiation, we additionally analyzed EVT-specific sFlt-1 (Fan et al., 2014) (Fig. 4 F). Well in line, EVT-, neutrophil-, and decBAM-associated factors were also significantly enriched in decB-CM with the exception of M-CSF, GM-CSF and CXCL1, the latter showing elevated but not significantly increased values (Fig. S6 A). Lastly, neutrophils secreted significantly higher amounts of S100A8, S100A9, and MMP9 (Fig. S6 B).

In summary, decBAMs differ substantially from decPAMs by a specific transcriptome and secretome.

### decBAMs induce Tregs and are suppressed in their APC-like function

Our IF analyses show a strong difference in decBAM and decPAM appearance, the latter exhibiting a more dendritic cell-like morphology (Fig. 5 A). Higher magnification clearly shows an enhanced spindle-shaped morphology and formation of pseudopodia in decPAMs, implying a more motile phenotype (Fig. 5 A). Interestingly, CD11c^lo^ decPAMs also showed a more prominent staining with antibody against HLA class II molecules (Fig. 5 B). Single channel and lower magnification pictures of CD14/CD11c and CD14/HLA-DRA co-stainings are shown in Fig. S7 A and B. Well in line, flow cytometry analysis reveals suppressed HLA-DR cell surface expression in decBAM cell isolates, which express high levels of CD11c (Fig. 5C). In contrast, decPAMs show high expression of HLA-DR and induced transcript levels of *HLA-DOA, HLA-DPA1* as well as *HLA-DRB1* and tended to show higher but not significantly elevated levels of *HLA-DRA* (Fig. S7 C). To test whether decPAMs indeed show pronounced APC-like functions we isolated decBAMs and decPAMs from the same donors and performed functional *in vitro* assays. Comparative video-based analysis of freshly isolated decidual macrophages indeed showed that decPAMs (Video 1) moved significantly faster than decBAMs (Video 2) when tracked in real time (Fig. 5, D and E). In addition, decPAMs also showed a significantly higher phagocytic activity when compared to decBAMs (Fig. 5, F and G). A recent study suggests that decidual macrophages are efficient inducers of Treg formation (Salvany-Celades et al., 2019).

**Figure 5.**
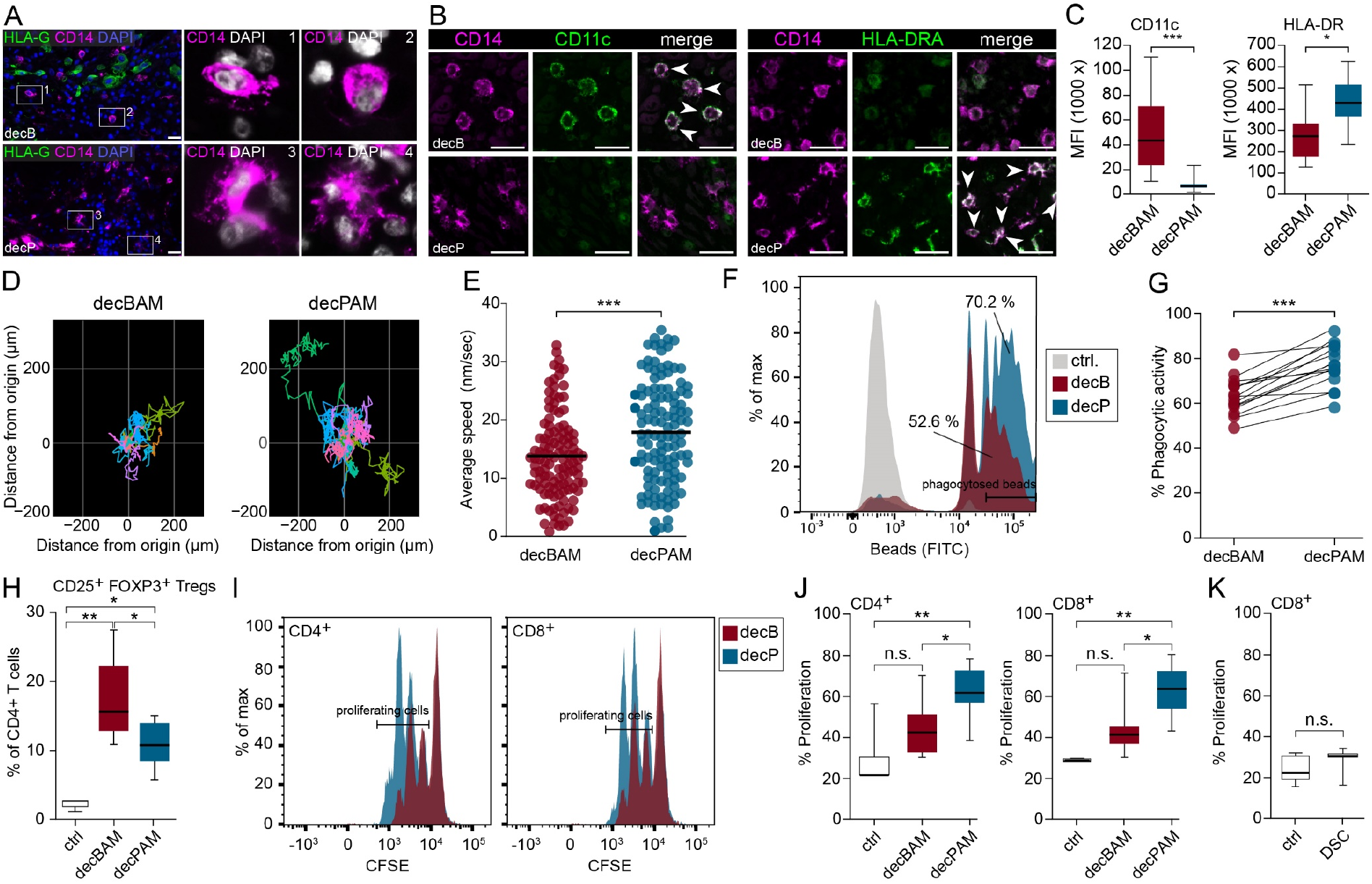
decBAMs show a pregnancy-tolerating phenotype by suppressed APC-like function and enhanced ability to induce Treg formation. **(A)** Representative IF staining of decB and decP tissue sections with antibodies against HLA-G (green) and CD14 (magenta). Zoomed insets on the right show appearance of CD14^+^ macrophages in areas invaded by HLA-G^+^ EVTs (1, 2) and those unaffected by placentation (3, 4). Representative images of *n* = 25 experiments. Scale bar, 20 µm. **(B)** IF co-stainings of first-trimester paraffin-embedded decB and decP tissue sections showing CD11c (green, left panel), HLA-DRA (green, right panel) and CD14 (magenta). White arrowheads indicate double positive cells. Representative images of *n* = 4 experiments. Scale bar, 50 µm **(C)** Median fluorescence intensity (MFI) of CD11c (n = 29 pairs= and HLA-DR (n = 8 pairs) was determined by flow cytometry in CD45^+^CD14^+^CD163^+^CD206^+^ decBAMs and decPAMs (n = 8 pairs) as indicated. The center line represents the median. Whiskers show the minimum and maximum of the data. P-values were calculated using paired Student’s t-test. **(D)** Representative cell tracks of isolated and short-term (24 h) cultivated decBAMs and decPAMs were generated by live-cell microscopy (capture rate = 12 frames/h). Cell tracks were color-coded for each monitored cell over 12 h. **(E)** Dots present average speed of captured individual cell tracks (n = 117) of decBAMs and decPAMs cultures (n = 8 pairs). 7 outliers were removed by Grubbs’ test. The center line represents the mean. P-value was generated using Student’s t-test. **(F)** Representative flow cytometric analysis of phagocytosed FITC-labeled beads by decBAMs and decPAMs after 2 h of incubation. **(G)** Dots represent the percentage of phagocytosed FITC-labeled beads of each individual experiment performed (n = 15 pairs). Patient-matched samples are indicated via connected dot plots. P-value was generated using a paired t-test. **(H)** Box plots show the potential of decBAMs and decPAMs to induce CD4^+^FOXP3^+^CD25^+^ Tregs from isolated CD3^+^ blood T cells (n = 5). P-value was generated using paired t-test. **(I)** Representative flow cytometry plots showing CFSE dilution in CD4^+^ and CD8^+^ isolated blood T cells stimulated with anti-CD3/CD28 in the presence of decBAMs (red histogram) or decPAMs (blue histogram). **(J)** Box plots represent the median level of the percentage of dividing CD4^+^ and CD8^+^ T cells (at least one cell division as defined by CSFE dilution) in the absence or presence of decBAMs and decPAMs (n = 8). Box plots represent median values ± SEM. Boxes in box plots represent upper and lower quartiles. The center line represents the median. Whiskers show the minimum and maximum of the data. DAPI was used to visualize nuclei. P-values were generated using one-way ANOVA with Tukey’s multiple comparison test. *, P ≤ 0.05; **, P ≤ 0.01; ***, P ≤ 0.001. **(K)** Box plots show the median levels of dividing CD8^+^ T cells in the absence or presence of DSCs (n = 3). P-values were calculated using Student’s t-test. decB, decidua basalis; decP, decidua parietalis; decBAM, decB-associated macrophages; decPAM, decP-associated macrophages; MFI, median fluorescence intensity; Tregs, regulatory T cells.

By studying this phenomenon, we found that induction of CD4^+^CD25^+^FOXP3^+^ Tregs is not a uniform feature of decidual macrophages since Treg induction was in particular executed by decBAMs (Fig. 5 H). To further test for dendritic cell-like function, we studied the potential of decidual macrophages to activate blood-derived T cells. To this end, we cultivated T cells from healthy donors with decBAMs or decPAMs in a ratio of 10:1. In line with our previous results, mainly decPAMs are able to trigger proliferation in both CD4^+^ helper T cells and CD8^+^ cytotoxic T cells (Fig. 5, I and J). We observed no induction of CD8^+^ T cells when blood-derived T cells were incubated with decidual stroma cells (DSCs), confirming that our assay is not disturbed by reactivity against allogeneic HLA class I molecules (Fig. 5 K).

Altogether, we show that decBAM are impaired in their ability to migrate, phagocytose, and promote T cell proliferation but are efficient inducers of Tregs.

### The secretome of EVTs induces a decBAM-like phenotype

Since our secretome screen of isolated EVTs revealed high secretion of immunomodulatory factors (Fig. 4 F) we assumed that EVTs can alter decidual macrophage activation and function. In order to avoid preconditioning by a decB environment likely under the control of EVTs, we decided to test this by studying EVT-dependent effects on either whole decP tissue explants or isolated decPAMs. To this end, we treated isolated decP tissue explants with EVT-conditioned media (EVT-CM) and investigated possible alterations in decPAMs upon treatment (Fig. 6 A). Immunofluorescence-based analysis indeed showed that addition of EVT-CM significantly increases CD44 expression of decPAMs *in situ* to a comparable level as detected in decB tissue explants (Fig. 6 B). Next, we exposed isolated decPAMs to 50% EVT-CM and found a significantly reduced motility and phagocytic activity (Fig. 6, C and D). EVTs secrete high levels of the immunomodulatory factors TGFβ1 and TGFβ2, well studied for their pleiotropic effects on macrophages, as well as high levels of PD-L1, which has recently been suggested to inhibit macrophage phagocytosis when expressed by tumor cells (Gordon et al., 2017). However, the suppressive activity of EVT-CM towards the phagocytic activity of decPAMs was not reversed by the addition of any of the blocking antibodies against TGFβ or PD-L1 (Fig. 6 E). Next, we investigated the effect of EVT-CM on the ability of decPAMs to interfere with T cell function. While neither EVT-CM alone nor in combination with decPAMs affected Treg differentiation (Fig. 6 F), we found a significantly reduced decPAM-dependent activation of CD4^+^ and CD8^+^ T cells upon treatment with EVT-CM (Fig. 6 G).

**Figure 6.**
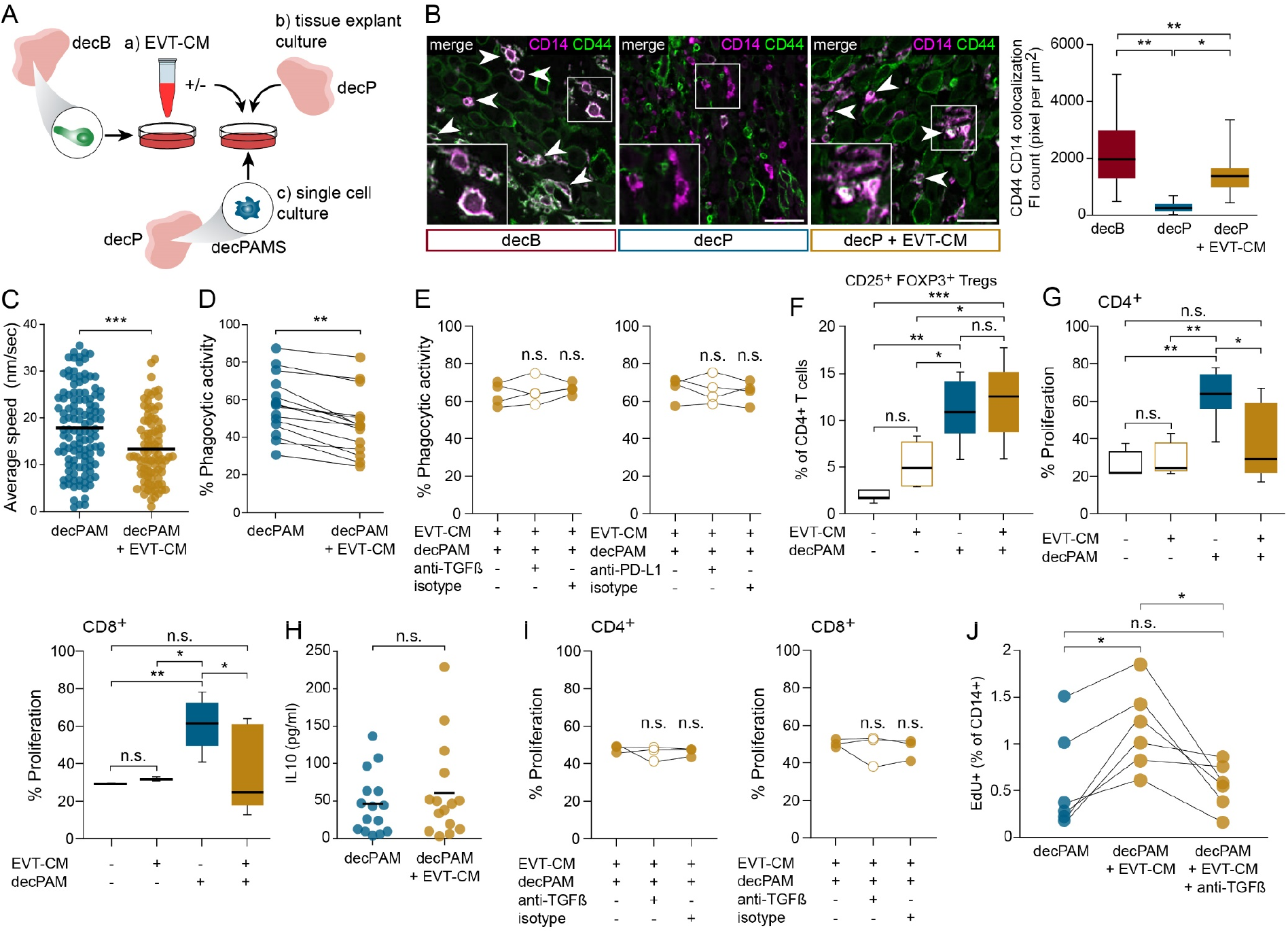
decPAMs are functionally reprogrammed by the secretome of EVTs. **(A)** Schematic drawing to illustrate the experimental set up. EVTs were isolated by magnetic beads conjugated with anti-HLA-G and cultivated for 20 h to generate EVT-CM (a) used to stimulate decP tissue explants (b) or FACS-isolated decPAMs (c). **(B)** Representative double IF stainings using antibodies against CD14 (magenta) and CD44 (green) investigating overlapping CD14 CD44 expression in unstimulated decB and decP tissue cultures and EVT-CM stimulated decP explant cultures. Examples of overlapping signals are marked by white arrow heads. Box plots indicate median value of white pixels per µm^2^ (generated by overlays of green and magenta colored CD14/CD44 co-stainings). P-values were generated using one-way ANOVA with Tukey’s multiple comparison test. Scale bars, 20 µm **(C)** Diagram shows average speed of each cell tracked (n = 90) in independent experiments using isolated decPAMs either without or with stimulation with 50% EVT-CM (n = 6 pairs). 7 outliers were removed by Grubbs’ test. The center line represents the mean. P-values was generated using a paired t-test. **(D)** Dot plot represents the percentage of phagocytosed FITC-labeled beads of each individual experiment performed (n = 14 pairs). Patient-matched samples are indicated via connected dot plots. P-value was generated using a paired t-test. **(E)** Phagocytosis assays were performed in the presence of EVT-CM alone, with isotype control or blocking antibodies for TGFβ or PD-L1. **(F)** Box plots show the potential of decPAMs and/or EVT-CM to induce CD4+FOXP3+CD25+ Tregs (n = 5). Box plots represent median values ± SEM. Boxes in box plots represent upper and lower quartiles. The center line represents the median. Whiskers show the minimum and maximum of the data. P-values were generated using One-way ANOVA with Tukey’s multiple comparison test. *, P ≤ 0.05; **, P ≤ 0.01, n.s.: not significant. **(G)** Box plots show the percentage of proliferative CD4^+^ or CD8^+^ T cells stimulated with anti-CD3/CD28 in the absence or presence of decPAMs and/or EVT-CM (n = 7). P-values were generated using one-way ANOVA with Tukey’s multiple comparison test. **(H)** Potential of short-term (20 h) cultivated decPAMs to secrete IL-10 (pg/ml) stimulated without or with EVT-CM (n = 15 pairs). **(I)** Box plots represent the percentage of dividing CD4^+^ and CD8^+^ T cells (at least one cell division as defined by CSFE dilution) stimulated with decPAMs and EVT-CM in the absence or presence of isotype control or a blocking antibody for TGFβ (n = 3). P-values were generated using one-way ANOVA with Tukey’s multiple comparison test. **(J)** Percentage of EdU^+^ nuclei in CD14^+^ macrophages of cultivated decP first-trimester tissue explants. Each dot represents the percentage for each tissue sample evaluated (n = 6 pairs). Patient-matched samples are indicated via connected dot plots. P-values were generated using Repeated-measures ANOVA with Tukey’s multiple comparison test. Boxes in box plots represent upper and lower quartiles. The center line represents the median. Whiskers show the minimum and maximum of the data. *, P ≤ 0.05; **, P ≤ 0.01; ***, P ≤ 0.001, n.s.: not significant. decB, decidua basalis; decP, decidua parietalis; decBAM, decB-associated macrophages; decPAM, decP-associated macrophages; EVT-CM, extravillous trophoblast-conditioned medium; Tregs, regulatory T cells.

Given that EVT-CM alone did not suppress CD4^+^ or CD8^+^ T cell proliferation (Fig. 6 G) we assumed an indirect effect for instance by EVT-CM-driven induction of IL-10, as previously shown for cancer-derived PD-L1 (Gordon *et al*., 2017) or by other TGFβ-driven effects on decPAMs. However, EVT-CM neither altered IL-10 secretion by cultivated decPAMs nor did TGFβ blockage alter the suppressive effect of EVT-CM on decPAM-induced T cell proliferation (Fig. 6, H and I). Lastly, we incubated decP explant tissues with EVT-CM and found significantly induced EdU incorporation into macrophages when compared to control treatment. In this setting, the addition of a neutralizing antibody against TGFβ reversed the growth-promoting effect of EVT-CM (Fig. 6 J).

In summary, these data show that EVTs have the capacity to trigger a pregnancy-tolerant phenotype in macrophages via their pleiotropically acting secreted factors.

## Discussion

In this study, we describe profound phenotypical and functional differences of tissue-resident macrophages in dependence on their localization within the human decidua. Moreover, the substantial difference between decBAMs and decPAMs in terms of transcripts, cellular proteome, and secretome also translated into different functional phenotypes. Finally, we show that EVT-CM is able to alter decPAMs into a phenotype functionally reminiscent of decBAMs. Surprisingly, we identified a significant increase in neutrophils and macrophages at the maternal-fetal interface. Except for a few studies in humans (Amsalem et al., 2014; Croxatto et al., 2016) and mice (Zhao et al., 2015), decidual infiltration with neutrophils is a poorly described phenomenon. To rule out incorrect tissue sampling and accidental contamination with blood-derived neutrophils, we established a thorough donor-matched tissue sampling strategy and, in addition, performed in-depth characterization of isolated neutrophils. In line with the negative L-selectin cell surface expression pattern identified on decB-derived neutrophils, we confirmed their interstitial presence using immunofluorescence and H&E stainings. Pregnancy is associated with accumulating numbers of blood granulocytic myeloid-derived suppressor cells (G-MDSCs) that can be distinguished from mature blood neutrophils by their different densities (Bronte et al., 2016). Indeed, we confirmed a significantly increased proportion of low-density neutrophils in the blood of pregnant women and found their accumulation in decB cell suspensions. Further characterization of decB-derived neutrophils revealed high levels of MMP-9 and the chemokines S100A8 and S100A9. While the latter two have recently been suggested to serve as markers for G-MDSCs (Bronte *et al*., 2016), MMP-9 production is a feature of tumor-associated neutrophils reported to trigger tissue remodeling and angiogenesis in this context (Ardi et al., 2007; Bekes et al., 2011). These data implicate that decidual neutrophils exert immunosuppressive function and may have additional functions in vascular remodeling.

We report a significant accumulation of CD14^+^ CD163^+^ CD206^+^ tissue macrophages in decB samples. There is a significant discrepancy in the literature regarding the frequency of macrophages in the human decidua that is presumably related to the differences in techniques, markers and tissues used to identify these cells (Bulmer et al., 1991; Haller et al., 1995; Helige et al., 2014; Repnik et al., 2008; Solders et al., 2017). Supportive evidence for the accumulation of macrophages at the maternal-fetal interface includes induced numbers of CD163^+^ cells in the vicinity of invading EVTs (Helige et al., 2014), increased levels of CD206^+^ and CD209^+^ cells in decB (Repnik *et al*., 2008), elevated distribution of CD45^+^ CD14^+^ cells in decB at term (Solders et al., 2017), and the accumulation of Ly6-C^hi^ monocytes/macrophages in the pregnant mouse uterus (Zhao et al., 2015). To account for these disparities, we established a gating strategy to identify true tissue macrophages in our optimized tissue collective. Indeed, CD14^+^CD163^+^CD206^+^ decidual macrophages revealed high expression of lineage-associated hallmark genes, typical morphology and other specific characeristics but failed to express prototypical monocyte-associated markers. Similar to our results obtained by determining CD14^+^ myeloid cells, we find a significant rise in CD14^+^CD163^+^CD206^+^ tissue macrophages in decB. We further sought to explore phenotypical and functional differences between decBAMs and decPAMs. Transcriptional profiling identified over 500 overexpressed DEGs in decBAMs compared to decPAMs, and subsequent enrichment analysis indicated them to be associated with neutrophil and myeloid cell activation. Well in line, isolated and short-term cultivated decBAMs actively secrete high levels of CXCL1, CXCL5, CXCL8, IL-10, and M-CSF. In particular, the latter two exert critical functions during pregnancy, for example, by inducing a decidual phenotype in monocyte-derived macrophages (Lindau et al., 2021; Svensson-Arvelund et al., 2015; Svensson et al., 2011) thereby acting in an autocrine self-perpetuating manner. Furthermore, reduced IL-10 blood levels are associated with human pregnancy disorders such as preeclampsia (Nath et al., 2020). While administration of IL-10 prevents naturally occurring fetal losses (Chaouat et al., 1995), its deletion can provoke spontaneous abortions in mice (Murphy et al., 2005). M-CSF has been shown to support placental function by affecting EVT invasion (Hamilton et al., 1998). In addition, we find that decBAMs overexpress specific cell surface markers including CD11c. While CD11c has been previously proposed as a marker for different macrophage populations in the decidua (Houser *et al*., 2011), our study is the first to localize CD11c^hi^ macrophages in decB. DEGs previously noticed to be expressed in isolated decidual CD11c^lo^ macrophages (Houser *et al*., 2011) largely clustered into decPAM-specific transcripts and vice versa DEGs found in to be expressed in CD11c^hi^ macrophages were overexpressed in decBAMs. Along these lines, our comparative computational approach demonstrated that our decBAM and decPAM populations share major similarities with dM1 and dM2 populations identified by scRNA-seq (Vento-Tormo *et al*., 2018), respectively. Along these lines, our study further complements previous reports identifying novel decidual macrophage populations (Houser *et al*., 2011; Svensson *et al*., 2011; Vento-Tormo *et al*., 2018) by specifying their localization to decB and decP.

We further revealed a significant difference in the cell surface expression of HLA class II molecules in isolated decBAMs and decPAMs. Functional assays revealed that decPAMs are efficient phagocytes, stimulate T cell proliferation and show a migratory phenotype. These data imply that decPAMs constitute a significant tissue-resident APC population, where they serve as patrolling cells to protect the maternal-fetal interface from invading pathogens. However, migratory APCs are central in priming T cell responses to peripheral tissue antigens (Allenspach et al., 2008; Itano et al., 2003) and thus pose a critical threat to human pregnancy by potentially provoking T cell responses to placental antigens. Indeed, we find that decBAMs are significantly impaired in their migratory and phagocytic activity and their potential to trigger T cell proliferation. In line with this, decBAMs induced higher numbers of CD4^+^CD25^+^FOXP3^+^ Tregs. This phenomenon is well-described for IL10- and TGFβ-secreting anti-inflammatory macrophages (Ruffell and Coussens, 2015) and has also been assigned to decidual macrophages (Salvany-Celades *et al*., 2019), however not in comparison to decPAMs. Accumulatively, our descriptive and functional data imply that decBAMs trigger a broad anti-inflammatory environment i T cell activation and are likely one of the main inducers of neutrophil influx.

Pioneering work in mice showed that so-called yolk sac-derived tissue-resident macrophages seed into tissues during embryogenesis and are replenished by local proliferation throughout life (Ginhoux and Guilliams, 2016; Palis et al., 1999). While the local renewal of tissue macrophages has been confirmed in various transplantation studies in the liver (Klein et al., 2007), brain (Ginhoux et al., 2010), lung (Bittmann et al., 2001; Byrne et al., 2020) and heart (Bajpai et al., 2018), a definitive proof for the existence of long-lived, tissue-resident macrophages in humans is missing. In our study we find that decBAMs show signs for DNA synthesis in an *ex vivo* assay and a higher proliferative capacity *in situ*, compared to decPAMs. Although we cannot exclude macrophage replenishment by monocyte extravasation, additional *ex vivo* assays suggest that decB-CMs do not exert higher potential to attract monocytes when compared to decP explant-CMs. Tissue macrophage proliferation at the maternal fetal interface has further been confirmed in Hofbauer cells, which likely replenish independently of the embryonic bone marrow (Thomas *et al*., 2021). However, tissue macrophage polarization greatly depends on extrinsic stimuli, regardless of ontogeny (Ginhoux and Guilliams, 2016). For instance, macrophage progenitors of different origins develop into fully functional alveolar macrophages (van de Laar et al., 2016) and peritoneal macrophage diversity is determined by tissue-derived signals (Okabe and Medzhitov, 2014). Therefore, we hypothesized that the phenotype of decBAMs depends on placenta-derived signals.

To test this, we incubated isolated decPAMs with CM of cultivated primary EVTs to mimic trophoblast-dependent effects on decidual macrophages unaffected by placentation in utero. Functional studies on viable tissue explants revealed that placenta-derived signals could override the tissue-specific phenotype of decPAMs. In this setting, the addition of EVT-CM led to a significant induction of proliferation and upregulation of CD44 expression, a marker of decBAMs. Paralleling the phenomenon observed in decP tissue explants, EVT-CM also changed the functionality of isolated decPAMs by suppressing their migratory and phagocytic activity. Similar observations have been made in cancer, where tumor cells selectively inhibit the phagocytosis of tumor-associated macrophages (TAMs) by avoiding the generation of tumor antigen-specific cytotoxic T cells (Pathria et al., 2019). Indeed, restoration of the phagocytic activity in TAMs led to cancer-specific priming of CD8^+^ T cells (Tseng et al., 2013). Similar to the reduced ability of decBAMs to trigger T cell proliferation, decPAM-mediated T cell activation was also significantly suppressed upon the addition of EVT-CM. These data show that EVT-derived factors play a central role in the extrinsic control of macrophage function. Neutralizing antibodies targeting TGFβ, PD-L1, or IL-10 did not affect the EVT-CM-dependent effects on decPAM, likely due to the pleiotropic effect of EVT-CM on decidual macrophages. Indeed, another study reported that the FOXP3+ Treg-inducing effect of EVTs was unaffected by the addition of neutralizing antibodies targeting CD3, HLA-C, HLA-G, ILT2, TGFβ, and PD-L1 (Salvany-Celades *et al*., 2019). The ineffectiveness of our attempts to block the effect of EVT-CM likely has several reasons -one of which may be trophoblast-specific glycosylation patterns. A recent report shows that specific sialylated glycans of placenta antigens mediate B cell suppression (Rizzuto et al., 2022). Indeed, glycosylation alters anti-body-mediated binding to PD-L1 and predicts therapy effectiveness. However, selective blockage of TGFβ reversed the growth-promoting effect of EVT-CM in decPAMs. This observation and our finding that isolated decBAMs secrete high levels of M-CSF imply a growth-inducing autocrine loop as well as extrinsic proliferative stimuli by invasive EVTs. Indeed, TGFβ1 has been shown to enhance macrophage proliferation via the induction of M-CSF (Celada and Maki, 1992).

The generation of an immunosuppressive environment is decisive for a successful pregnancy. However, our data impose that this phenomenon does not affect the entire decidua. In fact, we find that macrophages further away from the placentation site, referred to as decPAM in this study, show functionality of tissue-patrolling macrophages. In contrast, decBAMs residing at the maternal-fetal interface exhibit significantly less pronounced APC-like function. Our study further implies that this pregnancy tolerant phenotype is dictated by the placenta as EVTs can be used to modify the functionality of decPAMs towards a state reminiscent of decBAMs.

## Material and Methods

### Ethical considerations

All experiments and analyses were conducted in accordance with the Declaration of Helsinki as well as Austrian laws and guidelines. First trimester (6 – 10 weeks of gestation, n = 97) decidual and placental tissues were derived from women undergoing elective termination of pregnancy for nonmedical reasons. Pregnancy terminations were performed by surgical aspiration. The use of patient material was approved by the Ethics Committee of the Medical University of Vienna. Ethical approvals are annually renewed. The use of blood samples from healthy control women for migration assays of PBMCs was approved by the Regional Ethics Committee in Linköping, Sweden. Written informed consent was obtained from all subjects.

### Cell isolation

For decidual and endometrial cell isolation, tissue was cut into small pieces using a scalpel and digested using 2,5 µg/ml Collagenase IV (Worthington) and 25 mg/ml DNase I (Sigma-Aldrich) with the 37°C Multi-B program of a gentleMACS Dissociator (Miltenyi Biotec). The resulting single cell suspension was passed through a 70 µm cell strainer (Falcon). If necessary, red blood cells were lysed by incubating cells in a buffer containing 155 mM NH4Cl, 10 mM KHCO3, and 0.1 mM EDTA (pH 7.3) for 5 min, before staining for flow cytometry or sorting.

For flow cytometry and sorting, macrophages were defined as CD45^+^CD14^+^CD163^+^CD206^+^, monocytes were defined as CD45^+^CD14^+^CD163^-^CD206^-^, neutrophils were defined as CD45^+^CD66b^+^, and DSCs were defined as CD45^-^ HLA-G^-^. Dead cells were excluded using the viability dye Zombie Violet (BioLegend). For generation of conditioned media, sorted cells were seeded onto Nunc Cell-Culture plates (ThermoFisher) at a density of 1 million cells/ml in Decidua culture medium, consisting of DMEM/F-12 without phenol red (Gibco), supplemented with 10% FBS superior (Sigma Aldrich), 50 µg/ml gentamicin (Gibco) and ITS+ Premix (Corning), and cultured for 20 hours at 37°C. Cell culture supernatants were then centrifuged at 12000 g for 10 minutes at 4°C, snap frozen in liquid nitrogen, and stored at −80°C until further use.

PAMMs were isolated according to a published protocol (Thomas *et al*., 2021), with some adjustments. Briefly, villi were scraped with a scalpel and digested with 0.2% Trypsin (Invitrogen) and 12.5 mg/ml DNase I (Sigma-Aldrich) in HBSS-Mg/Ca-free medium (Sigma-Aldrich) at 37°C for 7 min. The resulting single cell suspension was passed through a 70 µm cell strainer (Falcon). If necessary, red blood cells were lysed by incubating cells in a buffer containing 155 mM NH4Cl, 10 mM KHCO3, and 0.1 mM EDTA (pH 7.3) for 5 min, before staining for FACS sorting. PAMMs were defined as CD45+HLA-DR+ cells.

Primary EVTs were isolated according to a previously published protocol (Haider et al., 2016). Briefly, villi were scraped with a scalpel and digested in two sequential digestion steps using 0.125% trypsin (Invitrogen) and 12.5 mg/ml DNase I (Sigma-Aldrich) in HBSS-Mg/Ca-free medium (Sigma-Aldrich). Each digest was performed at 37°C for 30 min without agitation. Subsequently, cells were filtered through a 100 μm cell strainer (Falcon). The cells were then layered on top of a Percoll gradient (10–70%; Pharmacia, Uppsala, Sweden) and centrifuged at 1,175 g at 4°C for 24 min without braking at the end of the run. Cells between 35% and 50% of the Percoll layer were collected. If necessary, red blood cells were lysed by incubating cells in a buffer containing 155 mM NH4Cl, 10 mM KHCO3, and 0.1 mM EDTA (pH 7.3) for 5 min. Remaining cells were then magnetically labeled with HLA-G – PE antibody and anti-PE microbeads and enriched for HLA-G+ EVTs by magnetic-activated cell sorting using an OctoMACS separator and MS columns (Miltenyi Biotec).

HLA-G+ EVTs were seeded on fibronectin-coated tissue culture plates (Nunc Cell-Culture Plates, ThermoFisher) at a density of 1 million cells/ml EVT culture medium, consisting of DMEM/F-12, GlutaMAX (Gibco), supplemented with 10% FBS Superior (Sigma Aldrich), 50 µg/ml gentamicin (Gibco), and 0.5 µg/mL fungizone (Gibco), and left to differentiate for 1-3 days before collecting cell culture supernatants. The conditioned media were centrifuged at 12000 g for 10 minutes at 4°C, snap frozen in liquid nitrogen, and stored at −80°C until further use.

### Cell density

To determine the density of cells within decidual tissues, the tissue was weighed before digestion and the weight was correlated with the cellular output after digestion, determined by a CASY TT cell counter.

### Western Blotting

Decidual tissue samples (decB and decP) were selected according to our established tissue sampling protocol described in Fig. S1, A and B. Tissue homogenization, lysis and determination of protein concentration was performed as recently published (Windsperger *et al*., 2020). Briefly, protein concentration was evaluated using a Bradford protein assay (adjusted to 1µg / µl in 1% sodium dodecyl sulphate) and additionally corrected by the evaluation of protein band intensities using a stain-free acrylamide solution (TGX FastCast, Bio-Rad). Prior to blotting bands were visualized by an imaging system (ChemiDoc, Bio-Rad) and band intensities were determined by Image Lab 6.0.1 (Bio-Rad) by calculating the Adjusted Volume (Adj.Vol) of all protein bands visualized. Proteins were then blotted onto polyvinylidene fluoride membranes (GE Healthcare, Buckinghamshire, UK), blocked with skimmed milk and incubated with primary antibodies (Table S1) overnight at 4°C. Finally, blots were incubated with horseradish peroxidase-conjugated secondary antibodies (Table S1). Signals were visualized with ChemiDoc Imaging System (Bio-Rad).

### Flow cytometry

For analysis of immune cell distributions within decidual tissues, cells were isolated as described above and stained for flow cytometry using the antibodies listed in Table S1. Immune cells were defined as CD45^+^. Within the immune cell gate, neutrophils were defined as CD66b^+^, macrophages/monocytes were defined as CD66b^-^CD14^+^, dNK/uNK cells were defined as CD3^-^CD56^hi^, T cells were defined as CD3^+^, and B cells were defined as CD19^+^. All defined immune cell types were mutually exclusive. Measurements were performed using a FACSVerse (BD) and DxFLEX flow cytometer (Beckman Coulter).

To create the TSNE plot in Fig. 1 we used the CyTOF workflow as published by Nowicka et al. (Nowicka et al., 2017) and modified it accordingly for flow cytometry data.

### Immunofluorescence of decidual tissue sections

First trimester decidual tissues were fixed in 7.5% formaldehyde and embedded in paraffin (Merck). Sections (2.5 μm) were cut using a microtome (HM355; Microm) and deparaffinized by incubating for 10 minutes in Xylene, followed by 100%, 96%, and 70% ethanol, and deionized water. Antigens were retrieved using citrate buffer (pH 6.0, Sigma-Aldrich) in a KOS MicrowaveStation (Milestone SRL). When indicated sections were stained with Hematoxylin and Eosin using an automated slide stainer with (Tissue Tek Prisma Plus, Sakura Finetek). For staining, sections were blocked for 1 hour in TBS/T containing 5% normal goat serum (Cell Signaling Technology) and then incubated with primary antibodies listed in Table S1 overnight at 4°C. Subsequently, the slides were washed in TBS/T 3 times and then incubated with secondary antibodies (2 µg/ml) and DAPI (1 µg/ml, Roche) for 1 hour at room temperature. Images were acquired on a fluorescence microscope (Olympus BX50) using a Hamamatsu ORCA-sparc digital camera and the Olympus cellSens Standard software.

### Explant culture

Explants were generated by cutting decB and decP tissue into small pieces of approximately 1×1 cm. Explants were weighed and put into 4 ml of Decidua culture medium per 1 gram of tissue, then cultured overnight at 37°C. Of note, decidual explant cultures have been shown to maintenance high viability over six days *ex vivo*, in culture (Hazan et al., 2010). Conditioned media were centrifuged at 12000 g for 10 minutes at 4°C, snap frozen in liquid nitrogen and stored at −80°C until further use.

To assess the effect of EVT-CM on macrophages *in situ*, explants were cultured in 50% EVT-CM overnight, then fixed in 7.5% formaldehyde and processed for immunofluorescence as described above. For proliferation assays, 10 µM EdU (EdU-Click 488 kit, baseclick) was added to the explant culture and the EdU detection protocol was performed according to manufacturer’s user manual before blocking. Sections were co-stained with CD14, and macrophage EdU incorporation was assessed using the ImageJ software.

For blockage experiments, 1 µg/ml anti-human TGF-β antibody was added to the culture.

### Analysis of conditioned media

Conditioned media from decB and decP explants were analyzed by the proximity extension assay on the Olink proteomics platform (www.olink.com), using the Inflammation Panel. Samples were sent to and analyzed by the Clinical Biomarker Facility Unit at SciLifeLab in Uppsala, Sweden.

Conditioned media from decB and decP explants, as well as decBAMMs, decPAMMs, EVTs, and decB neutrophils, were analyzed using the magnetic bead-based immunoassays by Bio-Techne (Luminex Performance Assay TGF-β Premixed Kit and Luminex Discovery Assay with personalized panels). The samples were processed according to manufacturer’s user manual and measured on a Luminex 200 machine.

### RNA sequencing

After isolation of macrophages or monocytes (as described above), RNA was extracted using the SPLIT RNA Extraction Kit (Lexogen) and sent to Lexogen in Vienna for library preparation and sequencing (QuantSeq). Fastq files were aligned to the human reference genome GRCh38 with Gencode annotation using STAR aligner in 2-pass mode (Dobin et al., 2013). The count matrices were calculated with STAR/htseq-count. Differential gene expression was calculated using DESeq2 (Love et al., 2014), applying batch modelling where required. Genes with counts lower than 100 in 90% of samples were filtered out. LFC shrinkage with approximate posterior estimation for GLM coefficients was applied for downstream visualization (Zhu et al., 2019). For the PCA visualization, batch effects were removed with limma (Ritchie et al., 2015). We calculated Gene Ontology enrichment scores with the webtool g:Profiler and plotted the data in R (Ashburner et al., 2000; Gene Ontology, 2021; Reimand et al., 2007).

scRNA-seq data were downloaded from EMBL-EBI using the accession number E-MTAB-6701. Raw sequencing data were aligned to the reference genome GRCh38 using 10x Genomics Cell Ranger 6.1.2. Output files were analyzed and filtered with the R package Seurat 4.0.6 (Butler et al., 2018; Hao et al., 2021; Satija et al., 2015; Stuart et al., 2019). All cells with low overall gene expression or which showed high mitochondrial gene expression were filtered out. The data were log-normalized before entering the Seurat integration workflow. Cluster annotation was performed applying the list of cluster-specific genes published along with the original publication (Vento-Tormo *et al*., 2018). Violin plots were created with the Seurat Vlnplot function, using either single gene expression or module scores, calculated with the Seurat AddModuleScore function (Butler *et al*., 2018; Hao *et al*., 2021; Satija *et al*., 2015; Stuart *et al*., 2019).

### PBMC migration assay

Blood from healthy, non-pregnant female donors was collected in Na-heparin tubes and PBMCs were isolated using Lymphoprep (Axis Shield) according to manufacturer’s instruction. Conditioned media of decB and decP explants were pipetted into wells of a 24-well plate (Corning) at a concentration of 25% CM in Transwell medium consisting of Roswell Park Memorial Institute (RPMI) 1640 medium supplemented with 1 mg/ml human serum albumin (Octapharma), and 100 units/ml penicillin, 100 µg/ml streptomycin and 29.2 mg/ml L-glutamine (Gibco). Transwell inserts (6.5 mm inserts with 5.0 µm pores, Corning) were coated with 1.43 mg/ml Matrigel (Growth factor-reduced, Corning) for 1 hour at room temperature. Then, the inserts were placed into the wells containing the CM and the plate was pre-incubated for 1 hour at 37°C. PBMCs were seeded into the top part of the transwell insert (0.5 million cells in 100 µl Transwell medium) and the plate was incubated at 37°C for 4 hours. Migrated cells were collected by taking up the medium from the bottom part of the transwell and scraping any adherent cells off the bottom of the plate. Cells were then stained for flow cytometry and analyzed using a FACSCanto II flow cytometer (BD). TruCount tubes (BD) were used to determine the absolute numbers of migrated cells.

### Neutrophil migration assay

Neutrophil migration towards decB and decP supernatants was measured using 96-well trans well plates (Corning, 5 µm polycarbonate membranes). Filters were pre-soaked in buffer (PBS with Ca2+/Mg2+ supplemented with 0.1% BSA, 10 mM HEPES, and 10 mM glucose) for 1 hr at 37°C. Polymorphonuclear leukocytes (PMNLs) were isolated from healthy, non-pregnant donors as previously described (Valadez-Cosmes et al., 2021). Bottom wells were loaded with 100 µl supernatants (1:2 dilution with media) and upper wells with 100 µl PMNL (4×106 cell/ml). Cells were allowed to migrate for 1hr at 37°C and subsequently fixed with 150 µl of fixation solution (CellFIX, BD). Number of migrated neutrophils was measured at a BD FACSCanto II for 30 sec and separated from eosinophils based on their different auto-fluorescence in the V450 channel.

### Cell Tracking

For in vitro analysis of cell motility, macrophages were seeded at a density of 250.000 cells/ml Decidua culture medium and incubated in a Lionheart FX automated live cell imager at 37°C and 5% CO2. Time-lapse videos were created by taking a picture every 5 minutes for 24 hours, using the Gen5 3.08 software. All images between 12 and 24 hours of culture were used to track cells and analyze their speed with the CellTracker tool in MATLAB (http://celltracker.website). Cell starting coordinates were set to 0,0 and subsequent data points were newly calculated accordingly for visualization purposes. Cell tracks were visualized in R.

### Phagocytosis Assay

Decidual cells were isolated as described above and incubated in the presence of fluorescence-labeled latex beads (50 beads per cell; latex beads, 1 µm, carboxylate-modified polystyrene, fluorescent yellow-green, Merck) for 2 hours at 37°C. A negative control was incubated for 2 hours at 4°C to control for non-specific adhesion of beads to the cell surface. Then, cells were detached from the plate using Accutase (Gibco StemPro Accutase, Fisher Scientific) and stained for flow cytometry. Phagocytosis was assessed on a FACSVerse flow cytometer using the BD FACSuite software, by measuring the signal of the beads within the macrophage gate (using the negative control as a gate-setting control).

To measure the effect of EVT-CM on macrophage phagocytosis, the cells were cultured in 50% EVT-CM. For blockage experiments, 1 µg/ml anti-human TGF-β or 20 µg/ml anti-human PD-L1 antibody was added to the culture.

### T cell assays

T cells were isolated from PBMC of healthy, non-pregnant female volunteers using the Pan T Cell Isolation Kit (Miltenyi,130-096-535). To monitor proliferation, T cells were labeled with 5 µM carboxyfluorescein diacetate succinimidyl (CFSE Cell Division Tracker Kit, Biolegend). For Treg induction and T cell activation, flat-bottomed 96-well plates (Nunc Cell-Culture Plates, ThermoFisher) were coated with 1 µg/ml anti-CD3 and 2 µg/ml anti-CD28 antibodies (BioLegend) for 2 hours and seeded with 50.000 T cells/well. To measure the effect of macrophages and EVT-CM on T cell proliferation and Treg induction, 5.000 macrophages and/or 50% EVT-CM were added to the culture. For the blockage experiments, 1 µg/ml anti-human TGF-β antibody was added to the co-culture. After three days, T cell proliferation was determined by CFSE dilution and Treg differentiation was determined by surface staining of CD25 and intracellular staining of FOXP3 using the eBioscience Foxp3/Transcription Factor staining buffer set (Invitrogen). Data were collected using a FACSVerse flow cytometer using the BD FACSuite software and were analyzed with FlowJo 10.7.

### Accession codes and data availability

BioProject accession number for RNA-seq of isolated samples: GSE196917

## Acknowledgments

We thank all patients and staff at Gynmed Clinic (Vienna) for providing placental and decidual samples. We thank the FACS Core Unit at the St. Anna Children’s Cancer Research Institute (Vienna) and Boris Kovacic and Dieter Printz (St. Anna Children’s Cancer Research Institute Vienna) are greatly acknowledged for technical help and advices with flow cytometry-based analysis and sorting. Emir Hadzijusufovic (Division of Hematology and Hemostaseology, Department of Internal Medicine I, Medical University of Vienna) is acknowledged for technical help with cell staining. The authors would like to acknowledge support of the Clinical biomarker facility at SciLifeLab Sweden for providing assistance in protein analyses. We thank Benjamin Schuster (Vienna University of Technology) for his help with computational analysis. Thomas Weichhart (Center of Pathobiochemistry and Genetics, Institute of Medical Genetics, Medical University of Vienna, Vienna) is acknowledged for scientific discussion.

This work was supported by the Austrian Science Fund (P33485 to J.P.) and by the Austrian National Bank (17613ONB to J.P.).

## Author contributions

Conceptualization, J. Pollheimer, S. Vondra, A.L. Höbler, A.I. Lackner, R. Bauer, J. Kargl, J., G. Schabbauer, Ernerudh. Formal analysis, J. Pollheimer, S. Vondra, A.L. Höbler, A.I. Lackner, J. Raffetseder, Z.N. Mihalic, V. Kunihs, P. Haslinger, S. Haider, J. Kargl, H. Husslein. Funding acquisition, J. Pollheimer. Methodology, J. Pollheimer, S. Vondra, A.L. Höbler, A.I. Lackner, J. Raffetseder, A. Vogel, S. Haider, R. Oberle, M. Wahrmann, J. Ernerudh, J. Kargl. Intellectual input, J. Pollheimer, S. Vondra, A.L. Höbler, A.I. Lackner, J. Raffetseder, A. Vogel, L. Saleh, S. Haider, M. Wahrmann, R. Bauer, J. Kargl, H. Husslein, P. Latos, G. Schabbauer, M. Knöfler, J. Ernerudh. Supervision, J. Pollheimer. Visualization, J. Pollheimer, S. Vondra, A.L. Höbler, A.I. Lackner. Writing, J. Pollheimer, S. Vondra, A.L. Höbler, A.I. Lackner.

All authors discussed the manuscript.

## Supplementary Data

**Fig. S1** presents a schematic illustrating how decidual samples were processed for characterization, their cellular densities after enzymatic digestion, and the characterization of decidual neutrophils. **Fig. S3** shows representative flow cytometry gating strategies. Fig. S3 shows the characterization of tissue-resident macrophages in decB and decP samples. **Fig. S4** shows the data for chemoattractant assays performed with decB- and decP-CMs using PBMCs and low magnification IF stainings of proliferative decBAMs. **Fig. S5** represents PCA and gene set enrichment analysis of RNA-seq data, comparative computational analyses, and single channel IF stainings of CD14^+^ decidual macrophages double positive for HMOX1 or CD44. **Fig. S6** shows the secretome of decidual explants and neutrophils. **Fig. S7** shows single channel and lower magnification pictures of CD14/CD11c and CD14/HLA-DRA co-stainings and transcript levels of ITGAX and HLA class II molecules in decBAMs and decPAMs. **Data S1** presents the complete list of measured factors by the Olink proteomics platform (related to Fig. 1, D). **Table S1** provides a list of antibodies used in this study.

