## Supplement Figures for "The human placenta shapes the phenotype of decidual macrophages"

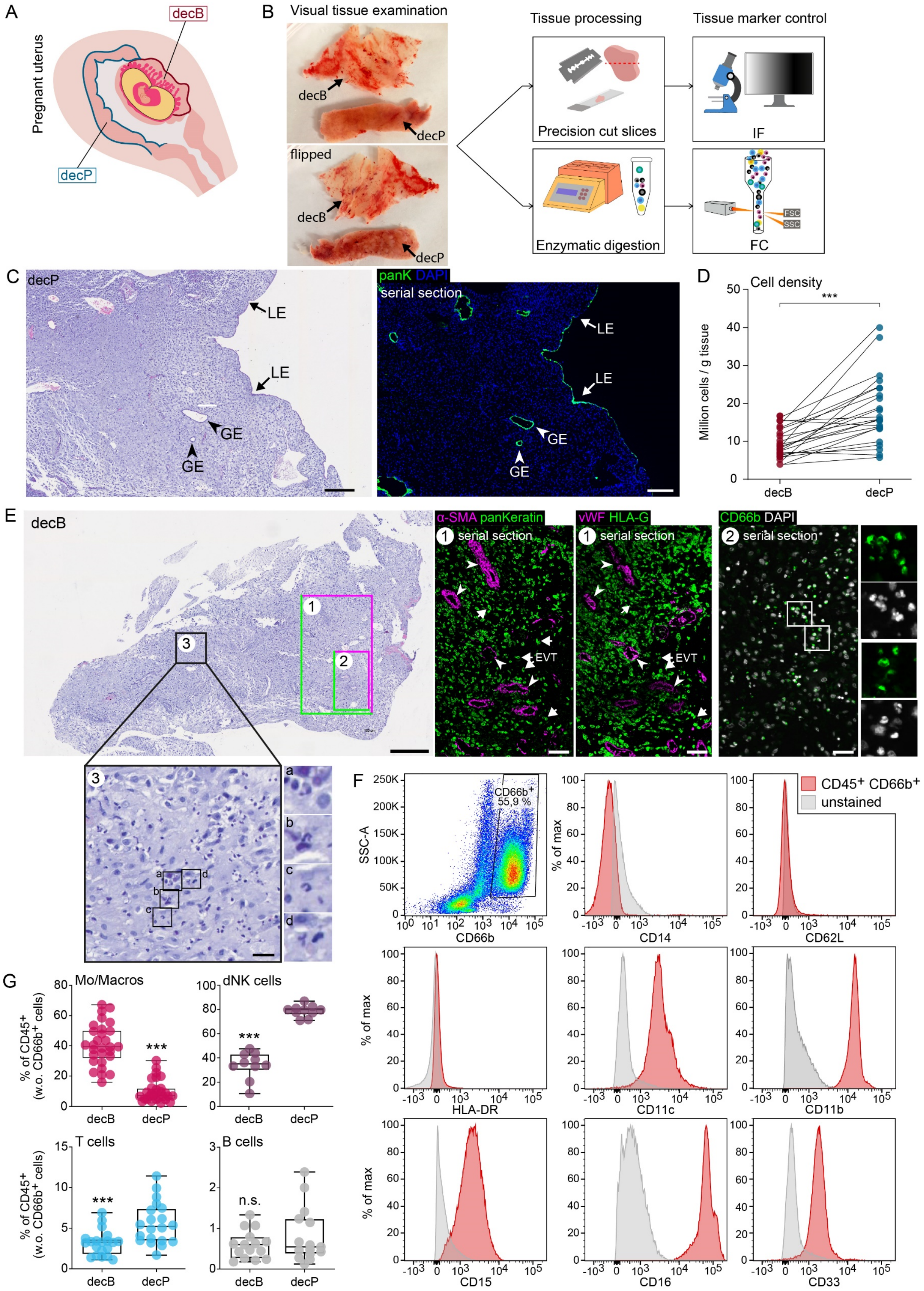


Figure S1 **Tissue processing, characterization and cell density of first-trimester decidual samples** **(A)** Schematic drawing of a pregnant uterus indicating decB and decP. **(B)** Schematic drawing of the tissue sampling strategy used in this study. Pictures to the left show representative photographs of the typical appearance of first-trimester decB and decP tissue samples. For control of proper sampling, each tissue is either processed for IF or flow cytometry analysis. **(C)** H&E staining of a representative decP tissue sample. Picture to the right shows a serial IF staining against panKeratin (panK) to visualize GE and LE as indicated by arrows and arrow heads, respectively. Scale bars, 200 µm. DAPI was used to visualize nuclei**.** **(D)** Dot plots represent cellular densities of processed decB and decP tissues (n = 24 pairs) determined by automated cell counting. Patient-matched samples are indicated via connected dot plots. P-value was generated using a paired t-test. **(E)** H&E staining of a representative decB tissue sample. Scale bar, 400 µm. Representative areas as indicated show IF stainings of serial sections (1) against α-SMA (magenta), panKeratin (green), vWF (magenta) or HLA-G (green) as indicated. Scale bars, 100 µm. White arrows indicate invasive EVTs and arrowheads point to blood vessels. The left outermost picture represents an IF-staining of a serial section (2) against CD66b (green). Scale bar, 50 µm. Zoomed images on the right show higher magnifications of CD66b+ (green) neutrophils with typical nuclear morphology. DAPI was used to visualize nuclei. Below of the H&E staining, a digitally zoomed higher magnification is shown of a selected area as indicated (3), representing neutrophil infiltrations. Scale bar, 20 µm. Neutrophils were evaluated based on their nuclear appearance. **(F)** The flow cytometry plot shows the $CD66b^+$ cell population of all $CD45^+$ cells from a cell suspension of decB. This population was analyzed for myeloid markers (CCD11b, CD11c, CD33), neutrophil-specific markers (CD16, CD15), macrophage-associated markers (CD14, HLA-DR), and a blood cell-specific marker (CD62L). Percentages indicate the frequency of the gated populations within total $CD66b^+$ cells. **(G)** Bar graphs show the percentages of Mo/Macros (n = 29 pairs), dNK cells (n = 10 pairs), T cells (n = 20 pairs) and B cells (n = 15 pairs) in the total CD45+ leukocyte population without $CD66b^+$ neutrophils as indicated. ***, P ≤ 0.001. α-SMA, alpha smooth muscle actin; H&E, Haematoxylin and eosin; decB, decidua basalis; decP, decidua parietalis; FC, flow cytometry; GE, glandular epithelium; IF, immunofluorescence; LE, luminal epithelium; vWF, Von Willebrand Factor.

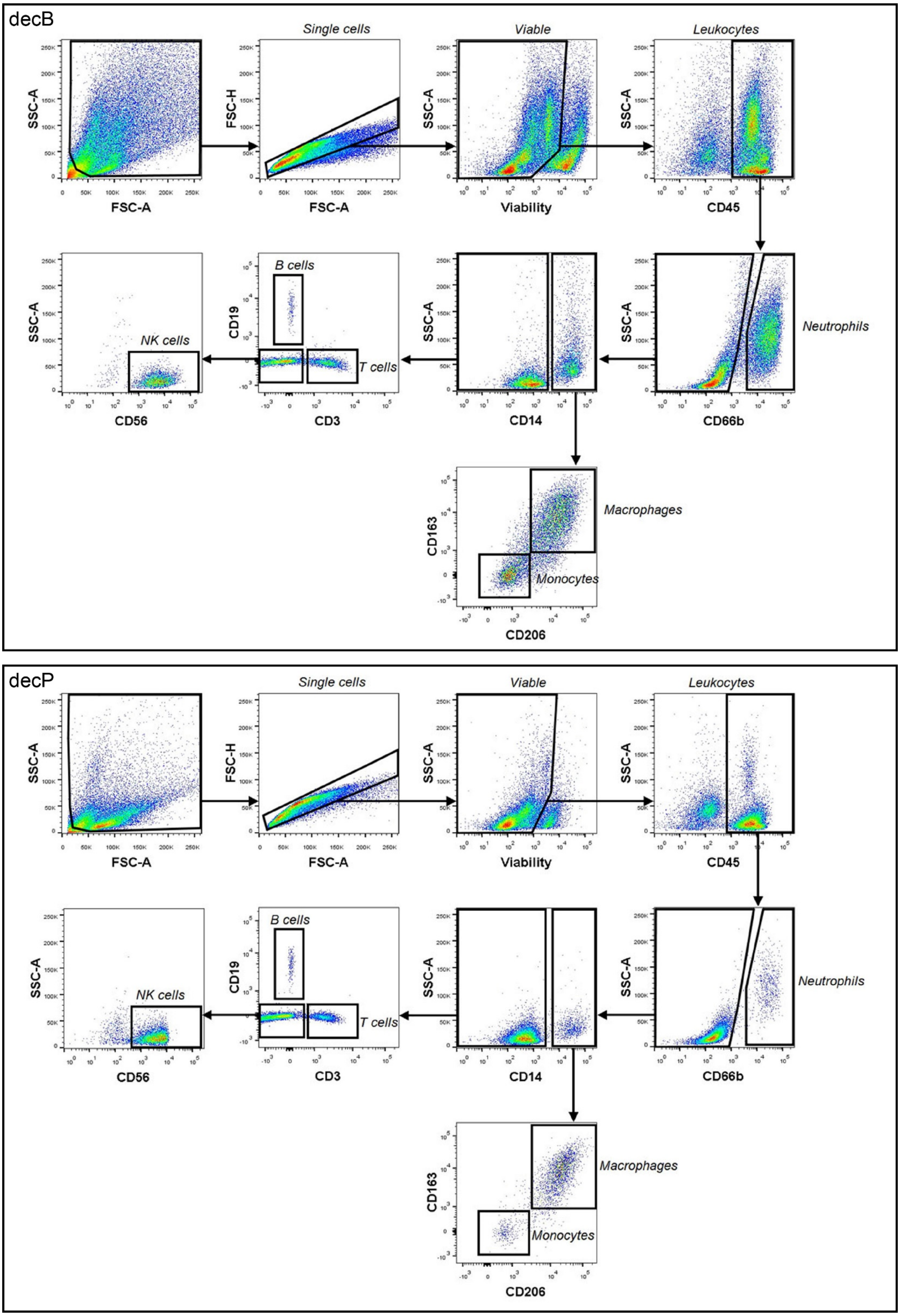


Figure S2 **Flow cytometry gating strategy for A) decidua basalis and B) decidua parietalis**

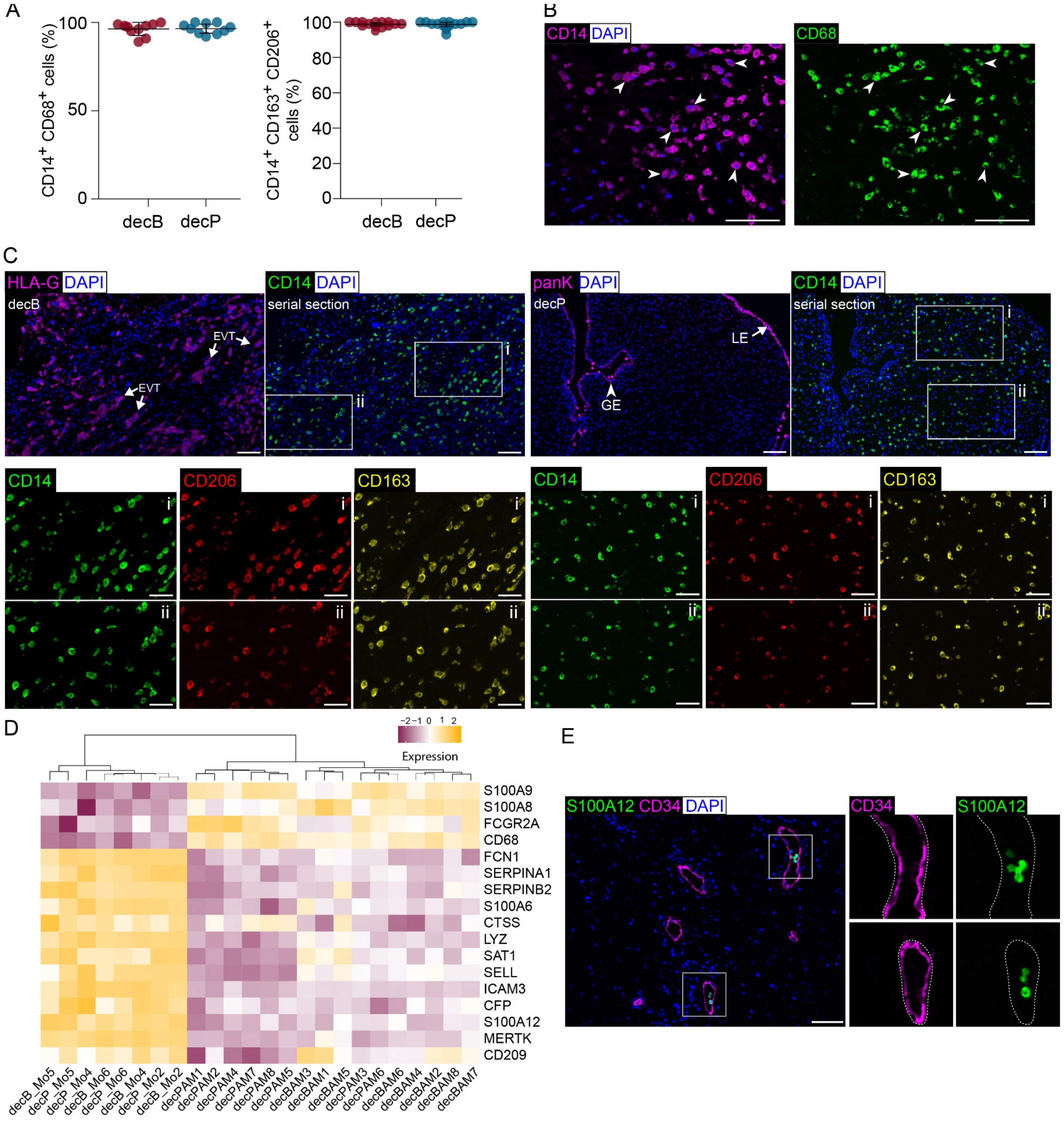


Figure S3 **Characterization of decidual macrophages and distribution of immune cells in non-pregnant endometrium and decidua. (A)** Percentages of CD14$^{+}$ CD68$^{+}$ (n = 10 pairs) and CD14$^{+}$ CD163$^{+}$ CD206$^{+}$ (n = 14 pairs) macrophages in patient-matched decB and decP tissues, as indicated. **(B)** Representative IF-staining for CD14$^{+}$CD68$^{+}$ cells in decidual tissues. Scale bars, 100 µm. **(C)** Representative IF staining using antibodies against HLA-G (magenta) or panK (magenta) of decB and decP samples, as indicated. Scale bars, 100 µm. Serial sections were stained with antibodies against CD14 (green), CD206 (red) and CD163 (yellow). Zoomed images below show higher magnification of areas indicated by white rectangles. Scale bars, 25 µm. **(C)** Heatmap showing expression of macrophage (pink) –and monocyte (yellow)-associated hallmark genes determined by bulk RNA-seq of isolated CD14$^{+}$, CD163$^{+}$, CD206$^{+}$ decBAMs and decPAMs as well as CD14$^{+}$, CD163$^{-}$, CD206$^{-}$ monocytes. **(D)** Representative immunofluorescence staining using antibodies against S100A12 (green) and CD34 (magenta). Zoomed images on the right show single channel expression of S100A12$^{+}$ monocytes in the lumen of CD34$^{+}$ vessels. Representative images of *n* = 3 experiments. Scale bar, 100 µm. decB, decidua basalis; decP, decidua parietalis; Endo, endometrium (non-pregnant); Mo, monocyte; decBAM, decB-associated macrophage; decPAM, decP-associated macrophage, dNK cells, decidua natural killer cells.

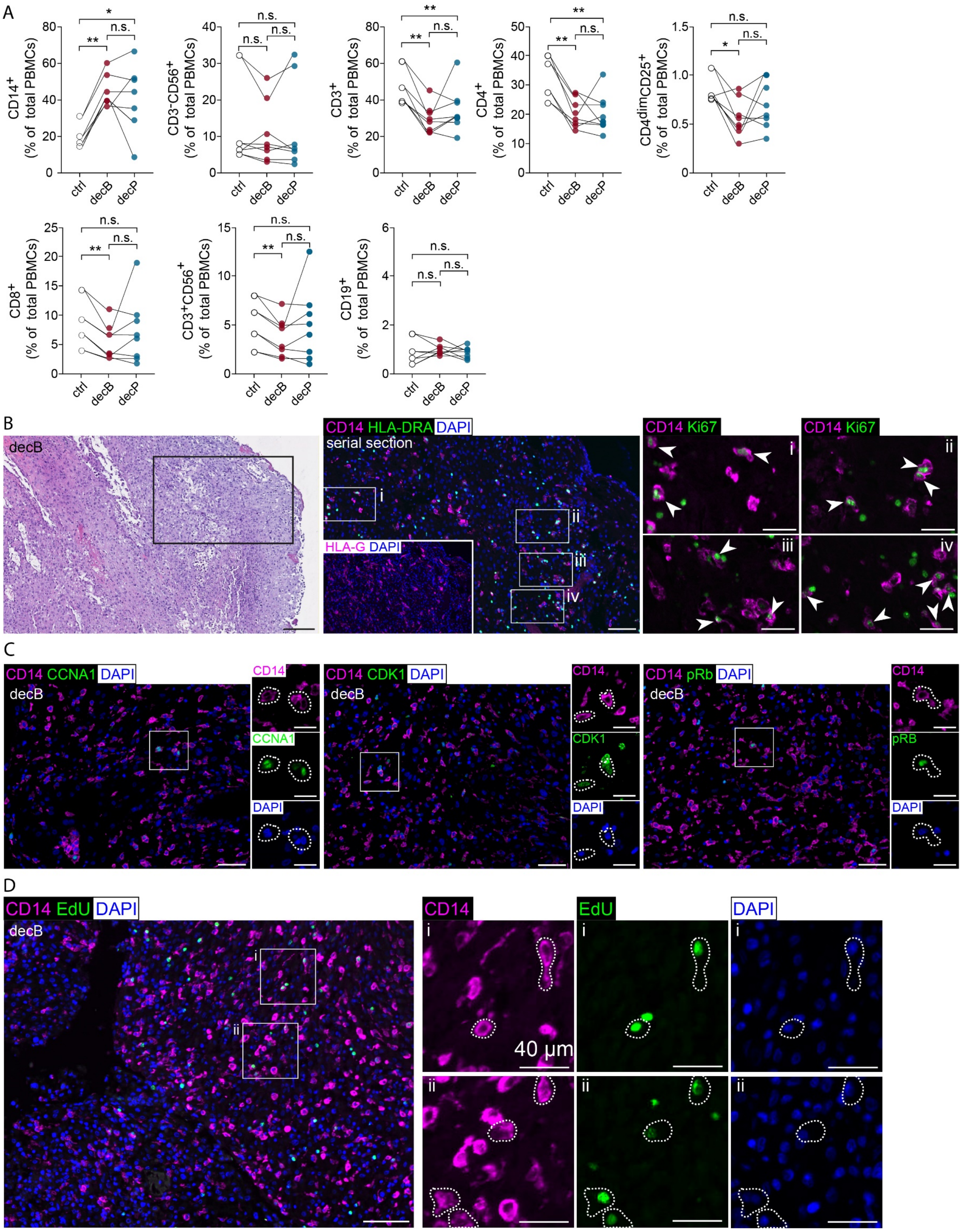


Figure S4 **Chemoattractant potential of decB- and decP-CM towards PBMCs and IF imagining of cell cycle maker expression in decB and decP tissue sections. (A)** Migratory response of PBMCs to culture medium containing CM of decB and decP explant cultures (1:4, n = 8). Decidua culture medium was used as a control. Percentages of individual blood immune cell types were analyzed by flow cytometry using antibodies against cell surface markers as depicted. P-values were generated using Repeated-measures one-way ANOVA with Tukey's multiple comparison test. *, P ≤ 0.05; **, P ≤ 0.01, n.s.: not significant. **(B)** H&E staining of a representative decB tissue sample. Scale bar, 250 µm. Picture to the right shows a serial IF staining of a higher magnification of an area as indicated by a black rectangle, stained against CD14 (panK) and ki67 (green). Double-positive cells are indicated by a white arrowhead. Scale bar, 100 µm. The inset in the left lower corner shows an IF staining against HLA-G to visualize EVTs. Zoomed insets to the right show higher magnification of areas indicated by white rectangles. Scale bars, 40 µm. DAPI was used to visualize nuclei. **(C)** Representative IF stainings of cell cycle markers (CCNA1, CDK1 and pRb; green) in decB tissue sections. Scale bars: 50 µm. Zoomed images to the right show higher magnification of areas indicated by white rectangles. Scale bars: 20 µm. DAPI was used to visualize nuclei. **(D)** Representative image of incorporated EdU (green) in $CD14^{+}$ (magenta) tissue macrophages of cultivated decB tissue explants. Scale bar: 100 µm. Zoomed images to the right show higher magnification of areas indicated by white rectangles. Scale bars: 40 µm. DAPI was used to visualize nuclei. decB, decidua basalis; decP, decidua parietalis.

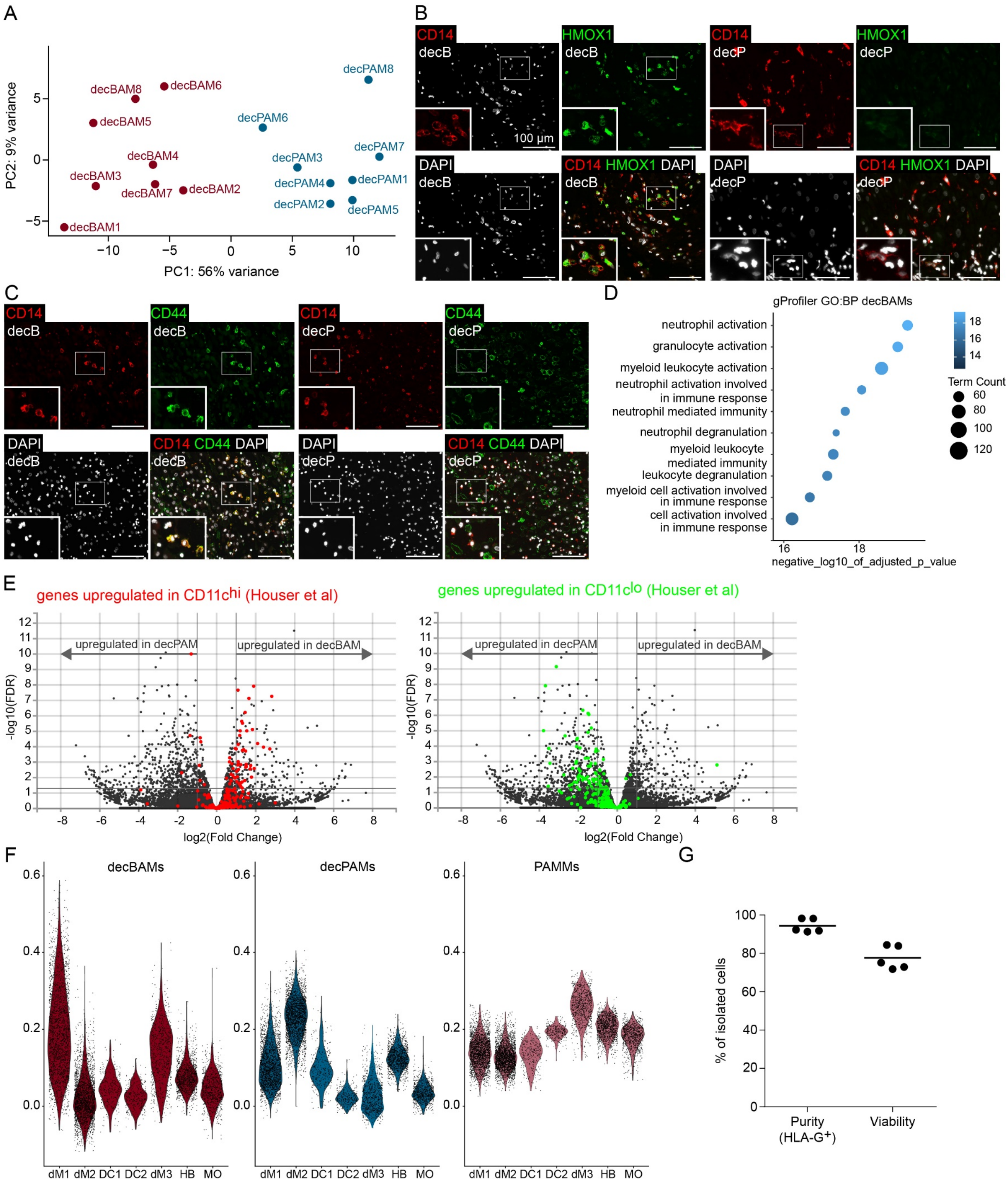

Figure S5 **Transcriptome and IF-guided analysis of decMs. (A)** Principal Component Analysis (PCA) of FACS-isolated, patient-matched decBAM and decPAM bulk RNA-seq expression profiles (n = 8 pairs). **(B-C)** Single channel stainings and lower magnifcations of IF-stainings presented in Fig. 4 B, as indicated. Insets on the left lower corner show higher magnifications of selected areas indicated by an white rectangle, Scale bars, 100 µm. **(D)** Gene set enrichment analysis of BP (biological processes) pathways using GO (gene ontology) datasets provided by g:Profiler. **(E)** Volcano plots depicting differentially expressed genes between decBAM and decPAM populations. Genes which were found to be differentially expressed between CD11c$^{hi}$ and CD11c$^{lo}$ macrophage populations (Houser et al.) were marked in red (upregulated in CD11c$^{hi}$) and green (upregulated in CD11c$^{lo}$). **(F** Violin plots showing expression of decBAMs (red), decPAMs (blue), and PAMMs (pink) signatures in selected scRNA-seq clusters (Vento-Tormo et al., 2018). **(G)** Assessment of purity (percentage of HLA-G$^{+}$ cells) and viability (percentage of zombie violet$^{-}$ cells) by flow cytometry. (n = 5).

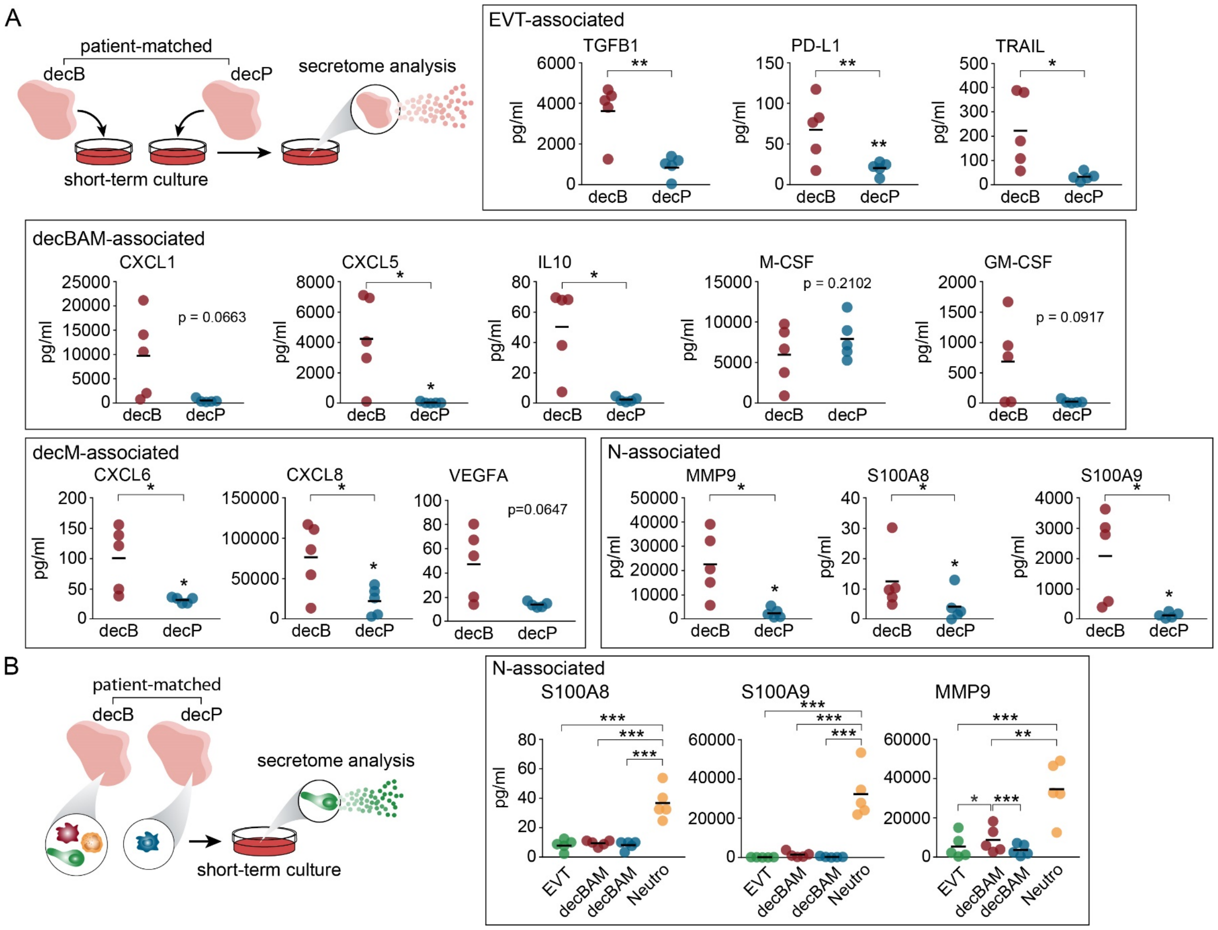


Figure S6 **Secretory potential of decidual explants and neutrophils. (A)** Secretory profile of cultivated (20 h), patient-matched decB and decP explants (n = 5 pairs) measured by bead-based multiplex immunoassays. Dots show measured levels in pg/ml. The center line represents the mean. **(B)** Secretory profile of FACS-sorted and cultivated (20 h) EVTs, decBAMs, decPAMS and Neutros, determined by bead-based multiplex immunoassays (n = 5 pairs). Measured levels (pg/ml) of growth factors, chemokines, and cytokines are shown as dots and the group mean is shown as centered line. The schematic drawing shown on the top illustrates the experimental set-up. P-values were generated using One-way ANOVA with Tukey's multiple comparison test. *, $P \leq 0.05$; **, $P \leq 0.01$; ***, $P \leq 0.001$. decB, decidua basalis; decP, decidua parietalis; decBAM, decB-associated macrophage; decPAM, decP-associated macrophage; PAMM, placenta-associated maternal macrophage; HB, Hofbauer cell; MO, monocyte; EVT, extravillous trophoblast; Neutro, neutrophil.

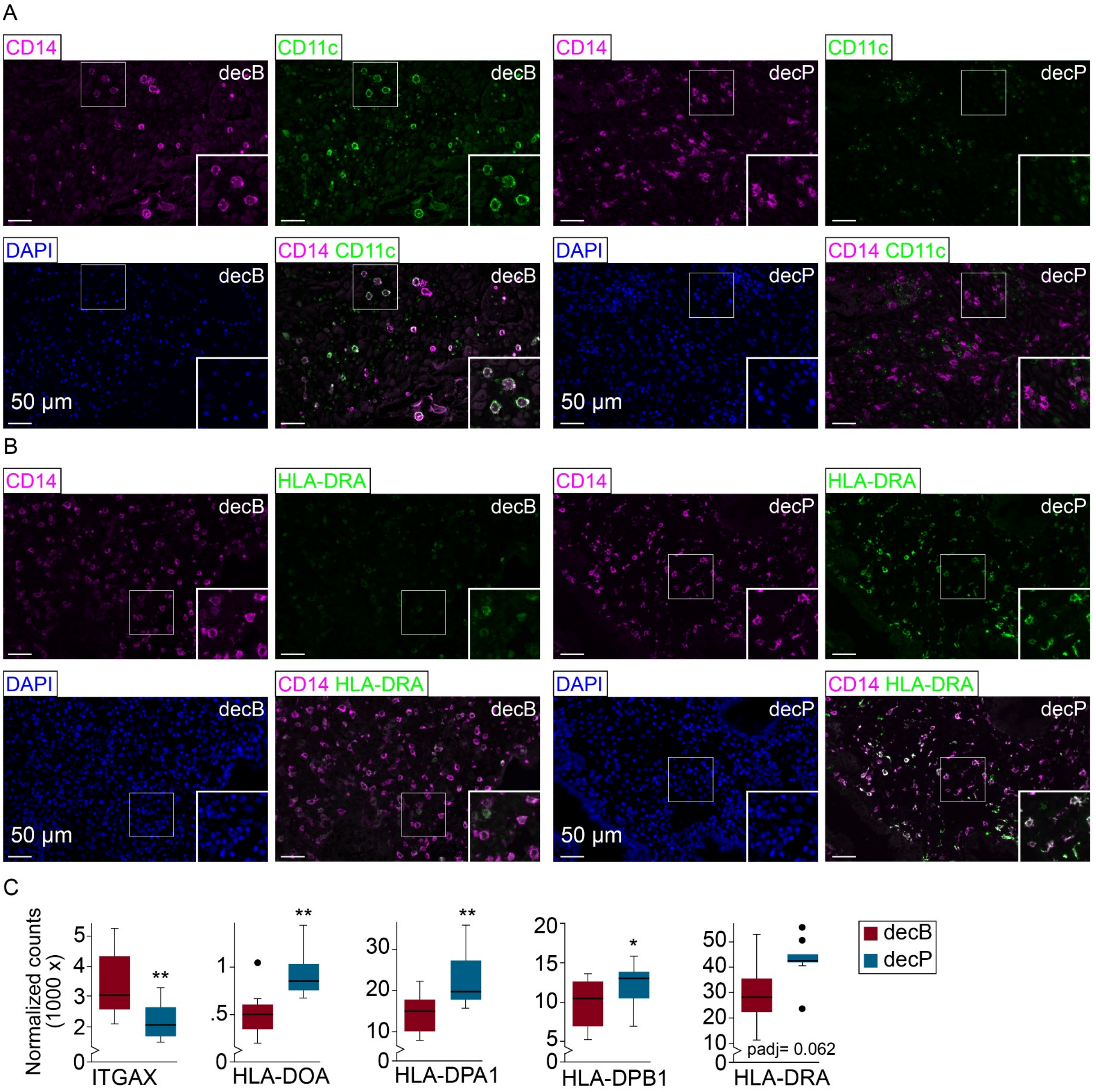


Figure S7**. MHC class II expression in decBAMs and decPAMs. (A - B)** Low magnification pictures of IF stainings presented in Fig. 5 B. Areas presented in Fig. 5 are indicated by a white rectangle. Scale bars, 50 µm. **(C)** Box plots show normalized transcript levels of ITGAX and HLA class II molecules in FACS-sorted CD14+CD163+CD206+ decBAMs and decPAMs analyzed by bulk RNA-seq (n = 8 pairs). Outliers are shown as dots and defined as data points outside 1.5 times the interquartile range above the upper quartile and below the lower quartile. P-values were calculated according to the recommended DESeq2 workflow.
