## Supplementary material for "The human placenta shapes the phenotype of decidual macrophages": Table S1

Short title: The placenta induces decidual macrophages

[§] These authors contributed equally to this work

### Abstract

During human pregnancy, placenta-derived extravillous trophoblasts (EVT) invade the decidua and communicate with maternal immune cells. The decidua can be distinguished into basalis (decB) and parietalis (decP), the latter being unaffected by placentation. By defining a novel gating strategy, we report accumulation of myeloid cells in decB. We identified a decidua basalis-associated macrophage (decBAM) population with a differential transcriptome and secretome when compared to decidua parietalis-associated macrophages (decPAMs). decBAMs are CD11c$^{hi}$ and efficient inducers of Tregs, proliferate *in situ* and secrete high levels of CXCL1, CXCL5, M-CSF, and IL-10. In contrast, decPAMs exert a dendritic cell-like, motile phenotype characterized by induced expression of HLA class II molecules, enhanced phagocytosis, and the ability to activate T cells. Strikingly, EVT-conditioned media are able to convert decPAMs into a decBAM phenotype. Cumulatively, these findings assign distinct macrophage phenotypes to decidual areas depending on placentation and further highlight a critical role for EVTs in the induction of pregnancy-tolerant macrophage polarization.


### Graphical Abstract

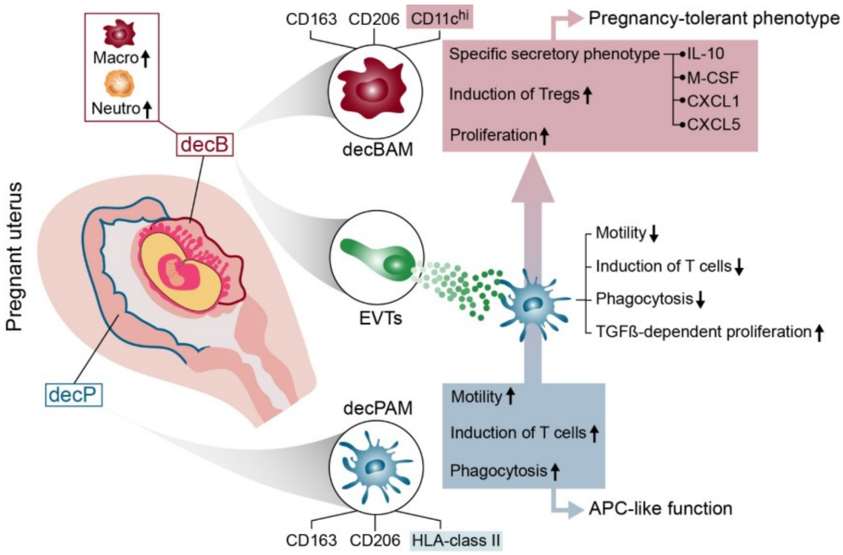


Highlights: In this study, we identified so far unrecognized, placenta-induced immune responses at the maternal-fetal interface. Altogether, we imply that placenta-derived trophoblasts induce a pregnancy-tolerant phenotype by suppressing antigen-presenting cell-like functions in maternal tissue macrophages.

### Results

#### Myeloid cells specifically accumulate at the maternal-fetal interface

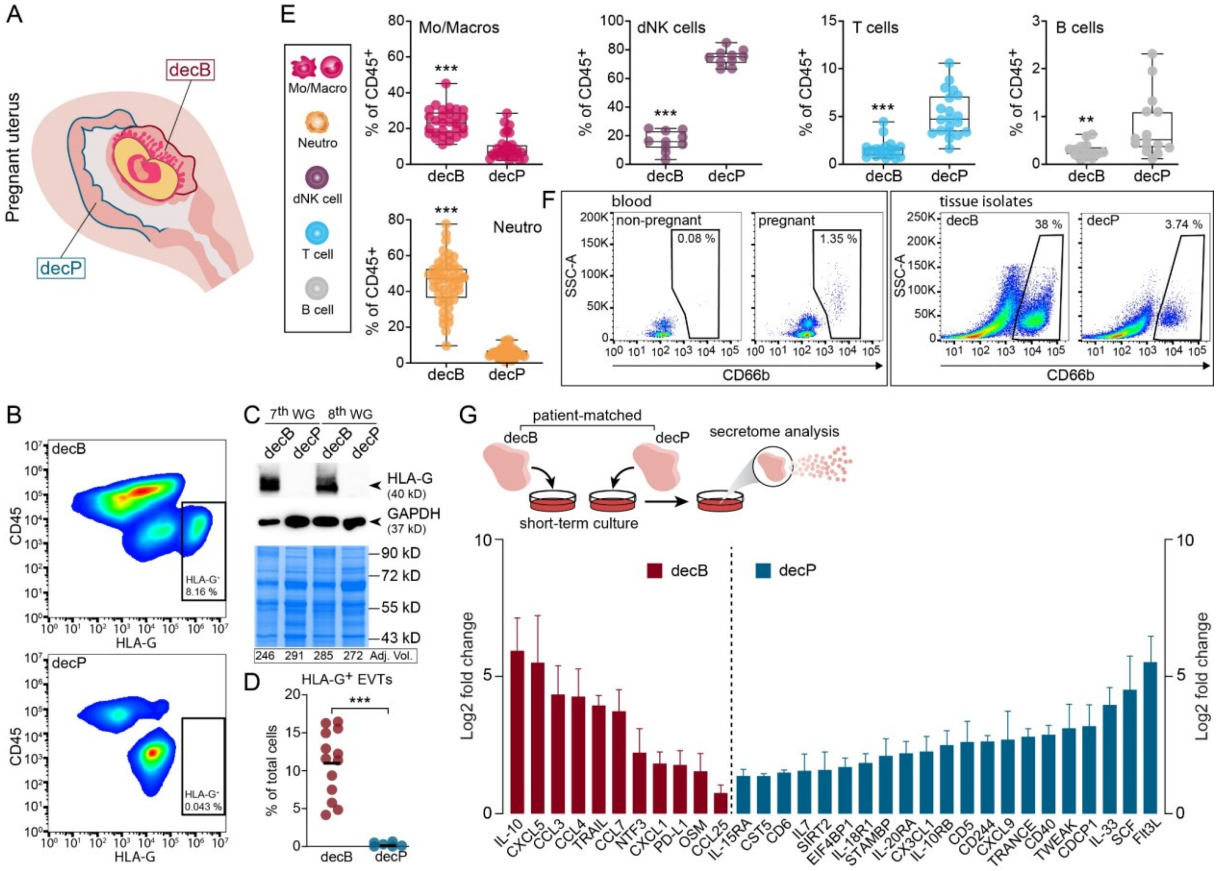


Figure 1.**Investigation of immune cell distributions in first-trimester decidual samples identifies myeloid cell accumulation in first-trimester decB tissues. (A)** Schematic drawing of a human pregnant uterus during the first trimester of pregnancy indicating decidua basalis (encircled by a red line, decB) and decidua parietalis (encircled by a blue line, decP). **(B)** Representative flow cytometry plots of decB and decP tissue cell isolates demonstrating distribution of $CD45^+$ immune cells and HLA-$G^+$ EVTs. The percentages of HLA-$G^+$ EVTs are indicated. **(C)** A representative western blot of two sets of donor-matched decB and decP protein lysates probed with antibodies against HLA-G is shown. The expression of GAPDH was determined to control for successful protein transfer and immunodetection. A stain-free system guaranteed equal total protein loading prior to blotting. Adjusted volumes (Adj. Vol.) of total band intensities are indicated underneath the stain-free gel images. **(D)** Quantification of HLA-$G^+$ EVTs in first-trimester decB (n = 14) and decP (n = 7) cell isolates, analyzed by Student's t-test. The center line represents the mean. **(E)** Bar graphs show the percentages of Mo/Macros (n = 31 pairs), dNK cells (n = 13 pairs), T cells (n = 21 pairs), B cells (n = 16 pairs) and Neutros (n = 53 pairs) of the total $CD45^+$ leukocyte population in donor-matched decB and decP tissue isolates. **(F)** Representative flow cytometry plots of isolated PBMCs from the blood of healthy non-pregnant and pregnant female donors (left panel) and donor-matched decB and decP tissue isolates (right panel). The gates represent the frequency of low-density $CD66b^+$ neutrophils of the $CD45^+$ PBMC fraction obtained by LymphoPrep®. **(G)** Summary data (log2 fold change, n = 4) for the significant differences in secreted factors from patient-matched decB and decP tissue explants after 20 h in culture measured by the proximity extension assay (www.Olink.com). The schematic drawing shown on the top illustrates the experimental set-up. Data are represented as mean + SD (standard deviation). Significances were calculated using multiple t-tests. **, $P \leq 0.01$. decGE, decidual glandular epithelium; EVT, extravillous trophoblast; Mo/Macro, monocytes, macrophages; Neutro, neutrophils; dNK cell, decidual natural killer cell; WG, week of gestation.

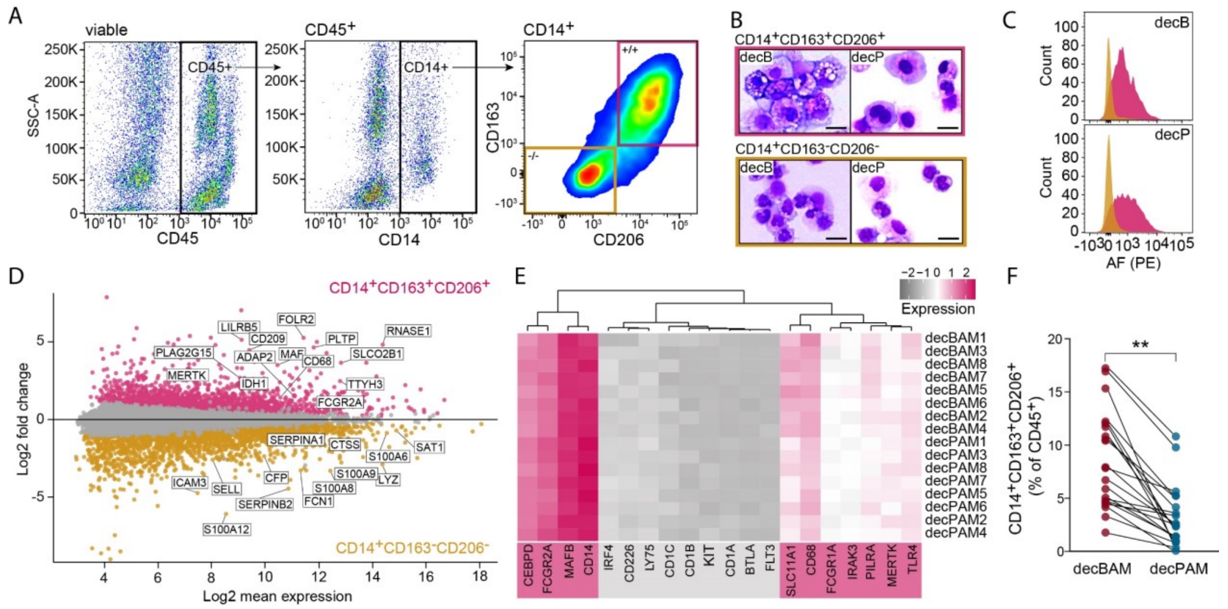


Figure 2. **Isolation and characterization of tissue-resident decidual macrophages (A)** Flow cytometry gating strategy used to identify tissue-resident decidual macrophages by a CD45$^{+}$CD14$^{+}$CD163$^{+}$CD206$^{+}$ expression profile. Pink gate, CD14$^{+}$CD163$^{+}$CD206$^{+}$ macrophages. Orange gate, CD14$^{+}$CD163$^{-}$CD206$^{-}$ monocytes. **(B)** Macrophage (pink rectangle) and monocytic (orange rectangle) populations from decB and decP were sorted and stained by Giemsa staining. Representative images of *n* = 4 experiments. Scale bar, 20 µm. **(C)** Representative flow cytometry plots identifying autofluorescence (AF) in isolated CD14$^{+}$CD163$^{+}$CD206$^{+}$ (pink histogram) and CD14$^{+}$CD163$^{-}$CD206$^{-}$ (orange histogram) cell isolates from decB and decP. **(D)** MA plot of differentially expressed genes (DEGs) from RNA-seq. Pink and orange dots represent significantly up- or down-regulated DEGs in isolated CD14$^{+}$CD163$^{+}$CD206$^{+}$ (n = 8 pairs) and CD14$^{+}$CD163$^{-}$CD206$^{-}$ (n = 4 pairs) cells from decB and decP tissues. **(E)** Heatmap showing expression of macrophage (underlaid with pink)- and DC (underlaid with grey)-associated hallmark genes in isolated decBAM and decPAM. **(F)** Dotplot demonstrating distribution of CD14$^{+}$CD163$^{+}$CD206$^{+}$ decBAMs and decPAMs in patient-matched first-trimester decB and decP samples (n = 21 pairs). Patient-matched samples are indicated via connected dot plots. P-value was generated using paired t-test. **, P ≤ 0.01. decB, decidua basalis; decP, decidua parietalis; decBAM, decB-associated macrophage; decPAM, decP-associated macrophage.

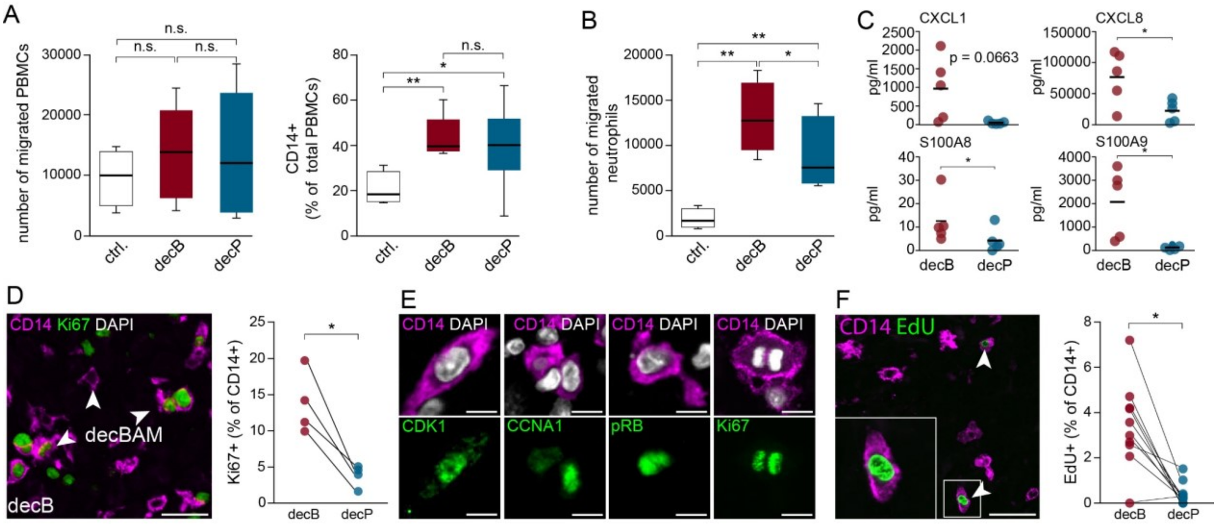


Figure 3. **decB-associated accumulation of myeloid cells is accomplished by chemoattractant factors and local proliferation of decBAMs (A)** Migratory response of PBMCs (left diagram) and CD14$^{+}$ blood cells (right diagram) to culture medium containing decB- or decP-CM (1:4, n = 8). Decidua culture medium was used as a control. P-values were generated using Repeated-measures ANOVA with Tukey's multiple comparison test. **(B)** Migratory response of blood neutrophils to culture medium without or with decB- and decP-CM (1:2, n = 4). Boxes in box plots represent upper and lower quartiles. The center line represents the median. Whiskers show the minimum and maximum of the data. P-values were generated using Repeated-measures ANOVA with Tukey's multiple comparison test. **(C)** Secretory profile of cultivated (20 hours), patient-matched decB and decP explants (n = 5 pairs) measured by bead-based multiplex immunoassays. Dots show measured levels in pg/ml. The center line represents the mean. P-values were generated using paired t-tests. **(D)** Representative IF staining of a first-trimester decB tissue section showing CD14 (magenta) and Ki67 co-staining (green). Quantification of four patient-matched decB and decP tissue samples is shown to the right by a dot plot. Each dot represents the percentage of Ki67$^{+}$CD14$^{+}$ double-positive cells for each tissue sample (n = 4 pairs). Scale bar, 20 µm. Patient-matched samples are shown in connected dot plots. P-value was generated using a paired t-test. **(E)** IF co-staining of first-trimester decB tissue sections showing cell cycle markers including CDK1, CCNA1, pRB in green and CD14 in magenta. The two images to the right upper and lower panel demonstrate a Ki67 expressing mitotic figure in a CD14$^{+}$ cell. DAPI (white) was used to visualize nuclei. Scale bars, 2 µm. **(F)** Representative IF staining of a cultivated decB explant tissue demonstrating EdU (green) incorporation into CD14$^{+}$ (magenta) macrophages, indicated by arrowheads. Inset picture shows a EdU+ macrophage at higher magnification. Quantification of EdU incorporation into CD14$^{+}$ macrophages of cultivated decB and decP first-trimester tissue explants. Each dot represents the percentage of CD14$^{+}$ EdU$^{+}$ cells for each tissue sample evaluated (n = 10 pairs). Patient-matched samples are indicated via connected dot plots. Scale bar, 20 µm. P-value was generated using a paired t-test. *, P ≤ 0.05; **, P ≤ 0.01, n.s.: not significant. decB, decidua basalis; decP, decidua parietalis.

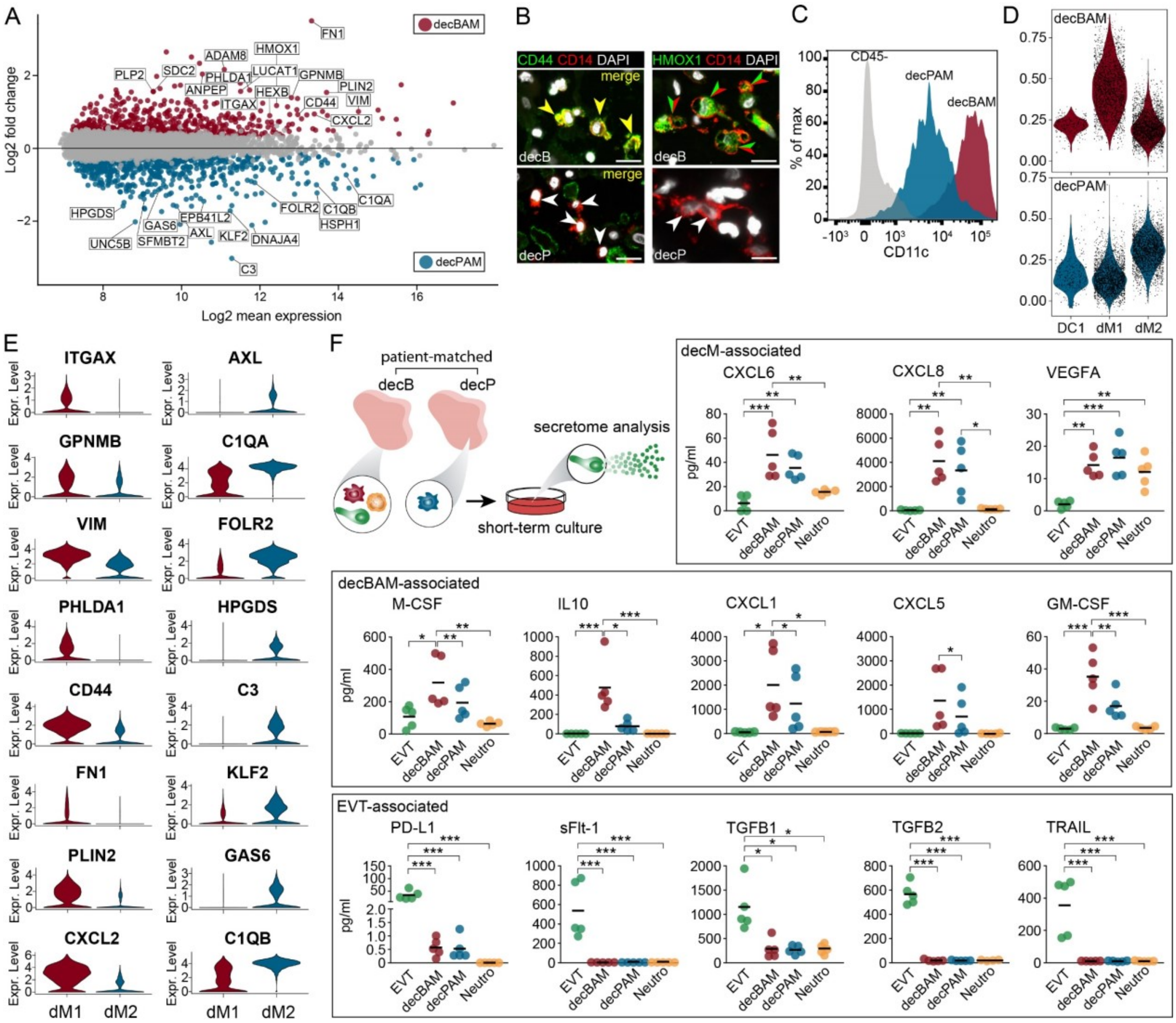


Figure 4. **First-trimester decBAMs and decPAMs differ in their transcriptional and secretory profile (A)** MA plot of DEGs from RNA-seq. Red and blue dots represent significantly up- or down-regulated DEGs in patient-matched isolated CD14$^+$CD163$^+$CD206$^+$ decBAMs and decPAMs (n = 8 pairs). **(B)** Representative IF co-staining of a first-trimester paraffin-embedded decB and decP tissue section showing HMOX1 and CD44 expression (green) in CD14$^+$ tissue macrophages (red). Arrowheads in yellow (left) or in green and red (right) indicate double positive cells. Scale bars, 20 µm. **(C)** Representative flow cytometric analysis of isolated decBAMs and decPAMs showing CD11c expression. **(D)** Violin plots showing expression of decBAMs (red) and decPAMs (blue) signatures in selected scRNA-seq clusters (Vento-Tormo et al., 2018). **(E)** Violin plots presenting expression of selected decBAM- and decPAM-associated gene transcripts in dM1 and dM2 scRNA-seq clusters (Vento-Tormo et al., 2018). **(F)** Secretory profile of magnetically sorted and cultivated (20 h) EVTs, decBAMs, decPAMS and neutrophils, determined by bead-based multiplex immunoassays (n = 5 pairs). Measured levels (pg/ml) of growth factors, chemokines and cytokines are shown as dots and the group mean is shown as centered line. The schematic drawing shown on the top illustrates the experimental set-up. P-values were generated using One-way ANOVA with Tukey's multiple comparison test. *, P ≤ 0.05; **, P ≤ 0.01; ***, P ≤ 0.001. decB, decidua basalis; decP, decidua parietalis; decM, decidual macrophage; decBAM, decB-associated macrophage; decPAM, decP-associated macrophage; EVT, extravillous trophoblast; Neutro, neutrophil.

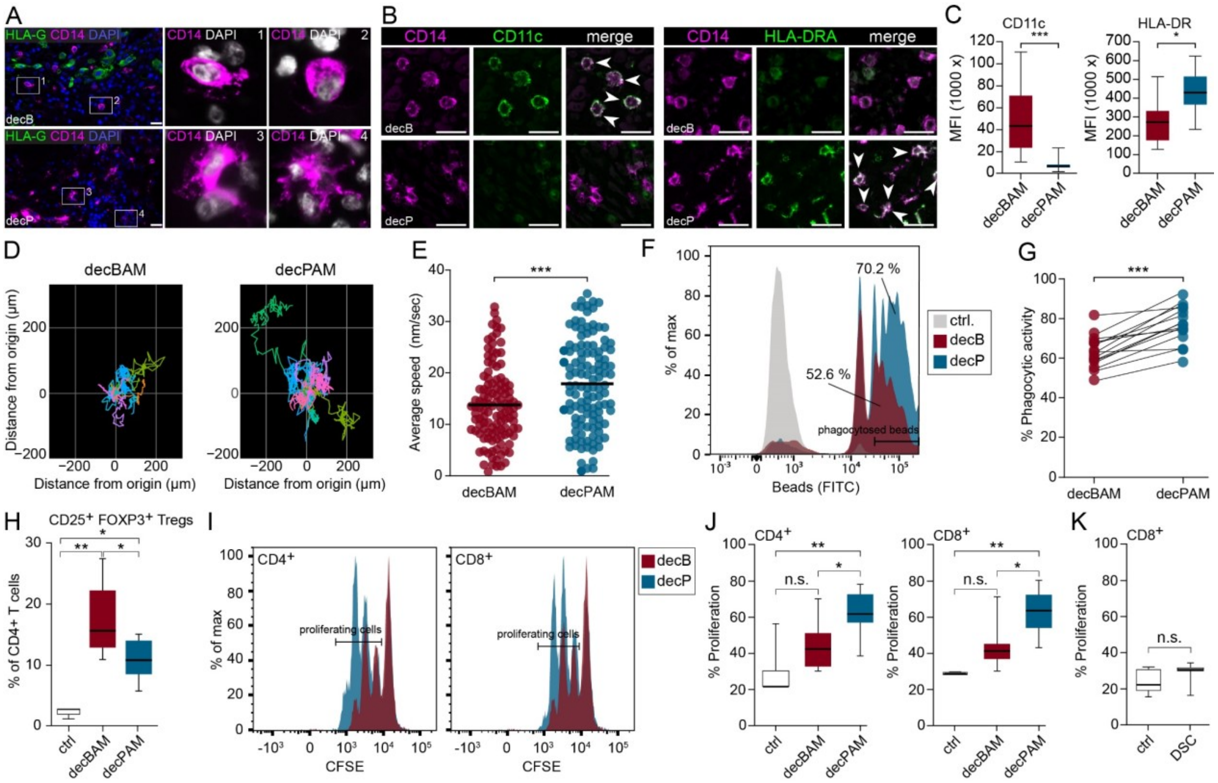


Figure 5. **decBAMs show a pregnancy-tolerating phenotype by suppressed APC-like function and enhanced ability to induce Treg formation. (A)** Representative IF staining of decB and decP tissue sections with antibodies against HLA-G (green) and CD14 (magenta). Zoomed insets on the right show appearance of CD14$^{+}$ macrophages in areas invaded by HLA-G$^{+}$ EVTs (1, 2) and those unaffected by placentation (3, 4). Representative images of *n* = 25 experiments. Scale bar, 20 µm. **(B)** IF co-stainings of first-trimester paraffin-embedded decB and decP tissue sections showing CD11c (green, left panel), HLA-DRA (green, right panel) and CD14 (magenta). White arrowheads indicate double positive cells. Representative images of *n* = 4 experiments. Scale bar, 50 µm **(C)** Median fluorescence intensity (MFI) of CD11c (n = 29 pairs= and HLA-DR (n = 8 pairs) was determined by flow cytometry in CD45$^{+}$CD14$^{+}$CD163$^{+}$CD206$^{+}$ decBAMs and decPAMs (n = 8 pairs) as indicated. The center line represents the median. Whiskers show the minimum and maximum of the data. P-values were calculated using paired Student´s t-test. **(D)** Representative cell tracks of isolated and short-term (24 h) cultivated decBAMs and decPAMs were generated by live-cell microscopy (capture rate = 12 frames/h). Cell tracks were color-coded for each monitored cell over 12 h. **(E)** Dots present average speed of captured individual cell tracks (n = 117) of decBAMs and decPAMs cultures (n = 8 pairs). 7 outliers were removed by Grubbs' test. The center line represents the mean. P-value was generated using Student’s t-test. **(F)** Representative flow cytometric analysis of phagocytosed FITC-labeled beads by decBAMs and decPAMs after 2 h of incubation. **(G)** Dots represent the percentage of phagocytosed FITC-labeled beads of each individual experiment performed (n = 15 pairs). Patient-matched samples are indicated via connected

dot plots. P-value was generated using a paired t-test. **(H)** Box plots show the potential of decBAMs and decPAMs to induce $CD4^+FOXP3^+CD25^+$ Tregs from isolated $CD3^+$ blood T cells (n = 5). P-value was generated using paired t-test. **(I)** Representative flow cytometry plots showing CFSE dilution in $CD4^+$ and $CD8^+$ isolated blood T cells stimulated with anti-CD3/CD28 in the presence of decBAMs (red histogram) or decPAMs (blue histogram). **(J)** Box plots represent the median level of the percentage of dividing $CD4^+$ and $CD8^+$ T cells (at least one cell division as defined by CSFE dilution) in the absence or presence of decBAMs and decPAMs (n = 8). Box plots represent median values ± SEM. Boxes in box plots represent upper and lower quartiles.

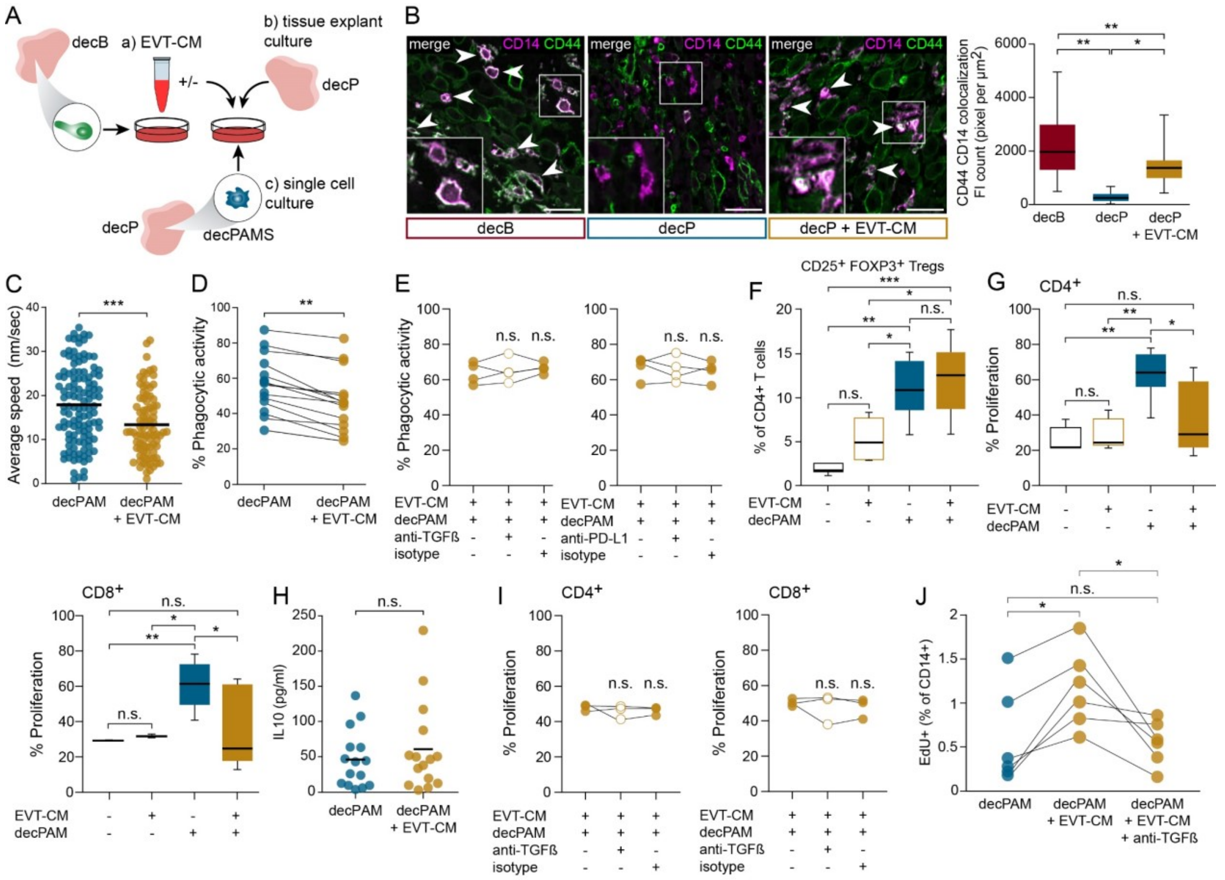


Figure 6. **decPAMs are functionally reprogrammed by the secretome of EVTs (A)** Schematic drawing to illustrate the experimental set up. EVTs were isolated by magnetic beads conjugated with anti-HLA-G and cultivated for 20 h to generate EVT-CM (a) used to stimulate decP tissue explants (b) or FACS-isolated decPAMs (c). **(B)** Representative double IF stainings using antibodies against CD14 (magenta) and CD44 (green) investigating overlapping CD14 CD44 expression in unstimulated decB and decP tissue cultures and EVT-CM stimulated decP explant cultures. Examples of overlapping signals are marked by white arrow heads. Box plots indicate median value of white pixels per $\mu m^2$ (generated by overlays of green and magenta colored CD14/CD44 co-stainings). P-values were generated using one-way ANOVA with Tukey's multiple comparison test. Scale bars, 20 µm **(C)** Diagram shows average speed of each cell tracked (n = 90) in independent experiments using isolated decPAMs either without or with stimulation with 50% EVT-CM (n = 6 pairs). 7 outliers were removed by Grubbs' test. The center line represents the mean. P-values was generated using a paired t-test. **(D)** Dot plot represents the

percentage of phagocytosed FITC-labeled beads of each individual experiment performed (n = 14 pairs). Patient-matched samples are indicated via connected dot plots. P-value was generated using a paired t-test. **(E)** Phagocytosis assays were performed in the presence of EVT-CM alone, with isotype control or blocking antibodies for TGFβ or PD-L1. **(F)** Box plots show the potential of decPAMs and/or EVT-CM to induce CD4+FOXP3+CD25+ Tregs (n = 5). Box plots represent median values ± SEM. Boxes in box plots represent upper and lower quartiles.
